# From Microstates to Macrostates in the Conformational Dynamics of GroEL: a Single-Molecule FRET Study

**DOI:** 10.1101/2023.03.23.533937

**Authors:** Demian G. Liebermann, Jakub Jungwirth, Inbal Riven, Yoav Barak, Dorit Levy, Amnon Horovitz, Gilad Haran

## Abstract

The chaperonin GroEL is a multi-subunit molecular machine that assists in protein folding in the *E. coli* cytosol. Past studies have shown that GroEL undergoes large allosteric conformational changes during its reaction cycle. However, a measurement of subunit dynamics and their relation to the allosteric cycle of GroEL has been missing. Here, we report single-molecule FRET measurements that directly probe the conformational transitions of one subunit within GroEL and its single-ring variant under equilibrium conditions. We find that four microstates span the conformational manifold of the protein and interconvert on the submillisecond time scale. A unique set of relative populations of these microstates, termed a macrostate, is obtained by varying solution conditions, e.g., adding different nucleotides or the co-chaperone GroES. Strikingly, ATP titration studies demonstrate that the partition between the apo and ATP-liganded conformational macrostates traces a sigmoidal response with a Hill coefficient similar to that obtained in bulk experiments of ATP hydrolysis, confirming the essential role of the observed dynamics in the function of GroEL.

**Significance Statement:** GroEL is a large protein-folding machine whose activity is accompanied by considerable conformational motions. Here, we use single-molecule FRET spectroscopy in combination with photon-by-photon statistical analysis to characterize the motions of a single GroEL subunit in real time and in the presence of ADP, ATP, and the co-chaperone GroES. Our results reveal transitions between four conformations on a timescale much faster than the functional cycle. We show that the motions of an individual subunit are directly coupled to the concerted allosteric mechanism of GroEL. This work, therefore, further demonstrates the impact of fast conformational dynamics on the biochemical function of molecular machines.

## Introduction

Molecular machines participate in multiple biological functions in the cell, such as the synthesis of biomolecules, cargo transportation, and DNA replication, to name a few. Such actions often involve converting available chemical energy, e.g., in the form of ATP molecules, into conformational changes with allosteric fine-tuning mechanisms (1). A prominent case study for investigating allosteric molecular machines is the oligomeric chaperonin GroEL, which assists in protein folding in the *E. coli* cytosol in an ATP-depended fashion (2-4).

First identified as a heat-shock protein (5, 6), GroEL is essential for the successful folding of certain proteins in the cell under either normal or stress conditions (7-10). It consists of 14 identical subunits arranged in two stacked heptameric rings, each forming a folding cavity that can accommodate a non-native protein substrate (11-14). The GroEL subunit can be segmented into three functional domains (**Figure 1 A**). The equatorial domain contains the nucleotide binding pocket (15, 16) and forms the inter-ring interface (13, 17). The apical domain is involved in the interaction with non-native protein substrates (18-20) and the co-chaperone GroES, a heptameric complex that occludes the folding cage with the substrate protein inside (12). Finally, the intermediate domain connects the equatorial and apical domains and plays an essential role in the activity of the protein (16).

**Figure 1:**
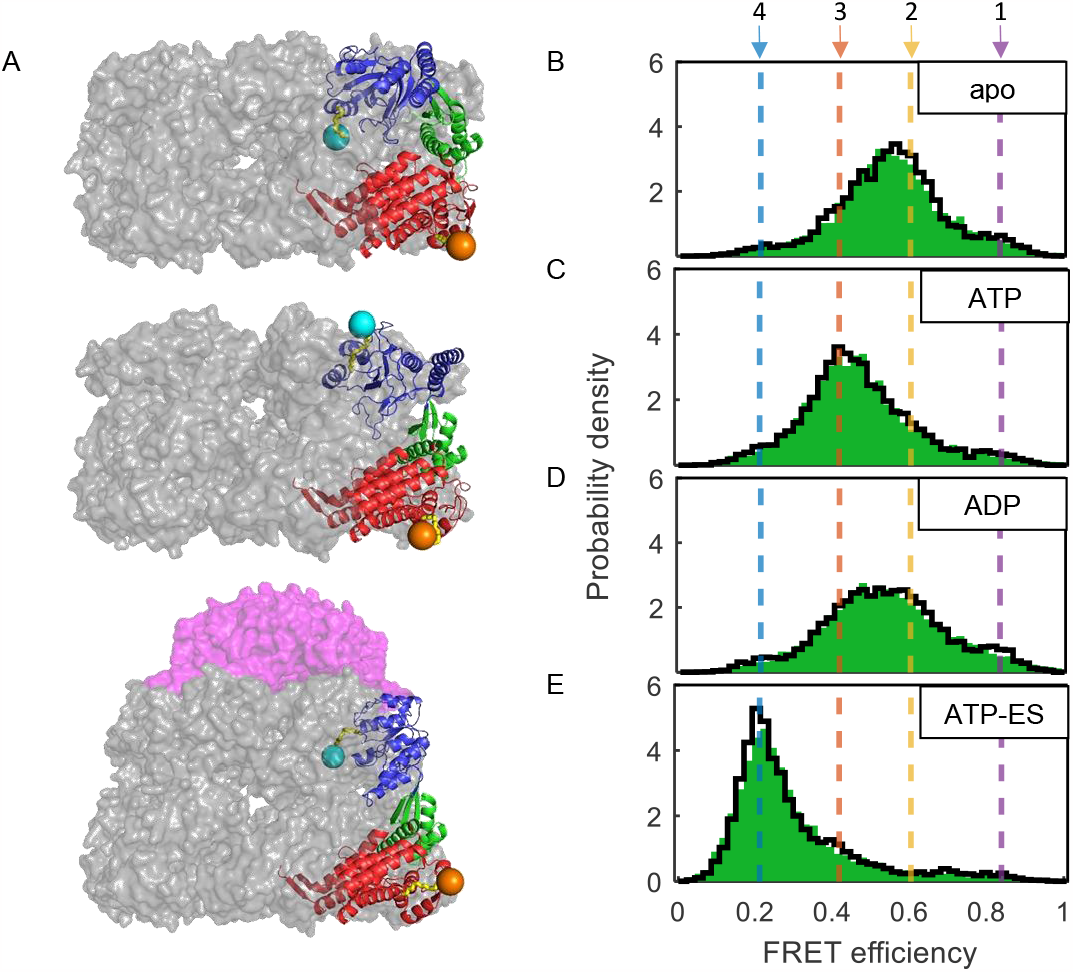
**A**) Structural models of the GroEL subunit within a single ring in different conformational states. From top to bottom: GroEL apo (PDB:1XCK (33)), GroEL bound to ATP (PDB:1AAR (35)), GroEL bound to GroES (PDB: 1AON (12)). Shown are the equatorial (red), intermediate (green), and apical (blue) domains of one GroEL subunit. GroES is shown in magenta. In each structure, a particular risFRET realization of the fluorescent labels is shown, with the linkers (yellow) and dye moieties at positions E255 (cyan) and D428 (orange). The inter-dye distance and expected FRET efficiency values from the risFRET simulations are *R* = 56 Å, *E* = 0.45 for the apo model, *R* = 67 Å, *E* = 0.25 for the ATP-bound model and *R* = 87 Å, *E* = 0.065 for the GroES-bound model. **B-E**) FRET efficiency histograms of SR1 255C/428C (green) under the following conditions: apo (**B**), 1 mM ATP (**C**), 1 mM ADP (**D**), and 1 mM ATP + GroES (**E**). The overlaid solid black lines are the “recolored” histograms generated using the model parameters from the global H^2^MM analysis (see text). The arrows and dashed lines mark the FRET efficiency values of the four microstates obtained from the analysis, which are 0.850 ± 0.002, 0.608 ± 0.005, 0.419 ± 0.006, and 0.213 ± 0.002 (mean ± standard error), respectively.

The complete reaction cycle of GroEL is governed by a seconds-long time window set by the hydrolysis of ATP and the consequent release of ADP/P_i_ (21), during which an encapsulated non-native protein substrate can start folding into its native state via several putative mechanisms (22-29). The allosteric transitions of GroEL with respect to ATP binding have been described by a nested cooperativity model, which assigns positive cooperativity between the seven subunits of one heptameric ring and negative cooperativity between the two stacked heptameric rings (30, 31). The intra-ring concerted transitions are described by the Monod-Wyman-Changeux (MWC) formalism (32), which posits an equilibrium between two allosteric states, T (tense) and R (relaxed), with low and high affinity towards ATP binding, respectively.

X-ray crystallography (11, 12, 14, 33, 34) and cryo-EM (13, 35-37) structures reveal that GroEL samples a rich conformational space with large domain motions during its reaction cycle. Upon ATP binding, the GroEL subunits transition from a closed to an open conformation in an upward motion of its apical domains. Further upward and clockwise twisting of the apical domains occurs when GroEL binds to the GroES heptamer. While most available structures were solved with an imposed C7 symmetry, in line with the concerted allosteric transition from the T to R allosteric states, a few studies have also shown stable asymmetric GroEL configurations, suggesting conformational heterogeneity among the protein subunits (34, 36).

Kinetic investigations of GroEL have identified several conformational processes spanning several orders of magnitude in time. Relatively slow events, such as substrate binding and release (38) or GroES exchange (16, 21) coupled to ATP hydrolysis, have been shown to occur within a few seconds. Other bulk transient kinetic experiments have indirectly probed conformational changes in the 10-200 milliseconds range (39-41), showing that the allosteric transitions triggered by ATP binding involve multiple kinetic phases.

Despite multiple studies on the allosteric conformational transitions of GroEL, a detailed, time-resolved description of domain motions within its subunits has not been presented to date, mainly due to the difficulties in collecting direct, highly resolved spatial and temporal data and the limitations of ensemble measurements. In this work, we address this deficiency and study the conformational dynamics of the GroEL subunit by measuring the local *equilibrium molecular motions* of a single subunit within the heptamer, using single-molecule FRET (smFRET) experiments on a single-ring variant of GroEL (42). Analyzing the data with a photon-by-photon statistical algorithm based on hidden Markov models (43), we find that the GroEL subunit transitions between four conformational microstates on a submillisecond time scale and that the relative population of these states depends on external conditions such as the presence and identity of nucleotides or GroES. Analysis of an ATP titration curve generated from single-molecule data provides direct evidence for the interrelation between the conformational states of the GroEL subunit and the allosteric transitions of the machine during its functional cycle.

## Results

### Designing a Labeled Variant of GroEL for smFRET Studies

In order to study the dynamics of a GroEL subunit using smFRET spectroscopy, we first searched for positions on the protein that, when labeled, would sample well its conformational changes. To that end, we used a program developed in the lab, risFRET (see Searching for Labeling Site Using risFRET in Methods), which simulates fluorescent probes onto PDB structural models of the whole GroEL complex and outputs a ranked list of potential FRET pairs. The ranking considers the expected FRET efficiency change between two specified conformations, the surface accessibility of the probes, and the conservation of residues (44). We eventually selected two labeling sites based on this analysis, E255 on the apical domain and D428 on the equatorial domain. **Figure 1 A** shows three structural models of the GroEL heptameric ring with a labeled subunit in the apo, ATP-bound, and GroES-bound states, each with an illustrative risFRET realization of the fluorescent dyes, AlexaFluor 488 and AlexaFluor 594, attached to the labeling positions with their linkers.

Throughout this work, we mainly used the single-ring variant of GroEL, SR1 (42), which, like the wild-type (WT) GroEL, exhibits ATPase activity with positive cooperativity, as shown in **Figure S2 A**. SR1 also interacts with the co-chaperone GroES in the presence of ATP. However, due to a lack of a negative allosteric signal from an opposite GroEL ring (16), the SR1-GroES complex remains stable for more than a hundred minutes (13, 24, 42, 45). This trait was utilized in our experiments. To detect FRET signals originating from only one GroEL subunit in the heptameric ring complex, we prepared SR1 constructs with a single double-labeled subunit using a disassembly-reassembly procedure (see Purification of GroEL Variants, Protein Labeling, and Reassembly Procedure in Methods and **Figure S1**). Briefly, purified and labeled 255C/428C monomers were mixed with unmodified SR1 subunits in a stoichiometric ratio of 1:100 in the presence of 4 M urea. After a short incubation, the mixture of monomers was dialyzed in a reassembly buffer, allowing the formation of SR1 rings. This mixing ratio guaranteed that ∼97% of the labeled SR1 complexes contained a single subunit labeled with the fluorescent dyes.

We performed a series of control experiments on the labeled and unlabeled SR1 constructs. An SR1 construct with seven unlabeled 255C/428C subunits demonstrated essentially the same ATPase activity as the WT SR1 variant, with a rate constant of 43 ± 2 min^-1^ and a cooperative response curve with a Hill coefficient of 2.0 ± 0.2 (**Figure S2 A-B**, see Steady-State Kinetic Assay in Methods). The integrity of various SR1 heptameric constructs used in this work was confirmed by comparing their migration pattern on a native gel and by native mass spectrometry measurements (**Figure S2 C-E**, see Denaturing and Native Gel Electrophoresis and Native Mass Spectrometry in Methods). The rotational freedom of the donor and acceptor dyes at each labeling site was verified with fluorescence anisotropy measurements on SR1 constructs with a single labeled protomer (see Fluorescence Anisotropy in Methods). Steady-state fluorescence anisotropy measurements of the labeled SR1 construct yielded values at 0.17-0.27 that did not change in the presence of ATP, ADP, and GroES (**Table S1**). Inspection of time-resolved anisotropy decay curves of each construct in the apo state showed a fast decay process with a correlation time of ∼1 ns (**Figure S3**), attributed to the rotation of the dyes, and a slow correlation time attributed to the rotation of the protein itself. These results indicate that the fluorescent labels are free to rotate on the protein, suggesting that changes in FRET efficiency observed in the experiments are only due to distance changes (46). We also validated the reported observations (47-49) that the three native cysteine residues of SR1 have a low labeling efficiency by attempting to label and prepare the WT variant in the same procedures as the single and double-labeled SR1 variants and conducting smFRET experiments (see Labeling Controls on Native Cysteine in Methods). Single-molecule stoichiometry histograms (**Figure S2 F**) demonstrated that the insertion of surface-exposed cysteine residues into SR1 reduced the labeling of the native cysteines significantly. Based on these results, we found the 255C/428C labeled SR1 construct as a suitable model to study the allosteric conformational changes of a single GroEL subunit within the heptameric ring.

### Single-Molecule FRET Resolves the Multi-State Conformational Space of the GroEL Subunit

We conducted pulsed-interleaved excitation smFRET experiments on diluted samples containing the labeled SR1 255C/428C construct without substrates (apo) or in the presence of either 1 mM ATP, 1 mM ADP, or 1 mM ATP with a stoichiometric excess of GroES relative to GroEL (see **Figure S4** and Single-Molecule Experiments in methods). In these measurements, labeled SR1 complexes freely diffused through a focused excitation laser beam, one molecule at a time, and emitted bursts of photons, also called photon trajectories (see below). The arrival times and the color of the detected photons in each burst were recorded, and the FRET efficiency and stoichiometry were calculated. For each condition, 2-3 measurements were conducted. **Figures 1 B-E** show FRET efficiency histograms (green) of selected measurement repeats (see **Figure S5** for corrected FRET efficiency histograms and **Figure S6** for uncorrected FRET efficiency of all measurement repeats). The rich conformational space of the GroEL subunit detected in all smFRET experiments is evident from the broad histograms, suggesting several distinct substrate-dependent conformational states. In the apo state (**Figure 1 B**), the FRET efficiency distribution exhibits a broad major peak centered at ∼0.55. In the presence of excess ATP (**Figure 1 C**), the FRET efficiency distribution is shifted to lower values, with the major peak now centered at ∼0.4, indicating that the apical domain is extended to a more open conformation. Interestingly, the addition of excess ADP (**Figure 1 D**) resulted in a broader FRET efficiency distribution centered between the apo and ATP peaks at ∼0.5. A dramatic shift in the FRET efficiency histogram was observed when the co-chaperone GroES was added in addition to excess ATP (**Figure 1 E**), with the peak value of the histogram now found at ∼0.2.

Since ATP bound to SR1 can hydrolyze to ADP, which may be released only slowly from the complex, it was possible that the recorded FRET efficiency distributions in the presence of ATP and ATP-GroES reflected a mixture of subunits bound to ATP and ADP. To examine this possibility, we conducted the same smFRET experiments in the presence of the ATP analog ADP-beryllium-fluoride (50) (ADP-BeFx, where x is the number of F-atoms, see Preparation of GroEL with ATP-Analog in Methods). This analog has been shown to mimic the ATP-bound state in GroEL, with the BeFx moiety acting as a stable substitution for the γ-phosphate of ATP (38, 51-53). The FRET efficiency histograms in the presence of ADP-BeFx, shown in **Figure S7**, are similar to those of the ATP-bound chaperone, indicating that ATP hydrolysis does not lead to the appearance of additional populations. Instead, these results confirm that the GroEL subunit can adopt several conformations even when indefinitely bound to ATP and GroES. In addition, these observations further emphasize the role of the γ-phosphate moiety in stabilizing the open, extended GroEL conformations. ADP alone does not seem to induce open conformations as effectively as ATP or ATP+GroES.

To verify that the observed conformational behavior of the SR1 heptameric ring reflects the behavior of the double-ring GroEL complex, we conducted smFRET experiments on the double-ring construct with the same labeling positions (prepared using the same reassembly procedure). The FRET efficiency histograms shown in **Figure S5 A-D** indicate that the two GroEL variants populate similar FRET efficiency states but with different amplitudes, as seen predominantly in the case of double-ring GroEL in the presence of excess ATP and GroES. These differences can be explained considering that in the smFRET experiment of the double-ring variant, half of the detected labeled subunits are in a GroES-bound (*cis)* ring, while the other half are in an unbound (*trans)* ring. This assertion was validated by forcing the double-ring variant to adopt the so-called symmetric football complex (53), where the two GroEL rings are bound to GroES (**Figure S5 E**). Since all 14 subunits now adopted the same GroES-bound state in this construct, we obtained a histogram that is more similar to the case of GroES-bound SR1. These observations strongly indicate that the conformational space sampled by a subunit in the heptameric ring is similar to that of a subunit in the double-ring GroEL complex.

To summarize, the labeled 255C/428C SR1 construct exhibits a conformational behavior that matches expectations based on prior knowledge. However, the existence of broad FRET efficiency distributions indicates that, during the different phases of its reaction cycle, the GroEL subunit transitions between several conformational states, with dynamics on the timescale of ∼0.5-1 milliseconds – the typical passage time of a molecule diffusing through the laser beam in the smFRET experiment.

### The GroEL Subunit Undergoes Fast Conformational Dynamics Between Four Conformational Microstates

To obtain a detailed description of the underlying conformational dynamics of the GroEL subunit, we performed a photon-by-photon hidden Markov model analysis of single-molecule photon trajectories using the program H^2^MM (43) (see H^2^MM Analysis in Methods). For each dataset, which consists of ∼6000-10000 photon trajectories, the H^2^MM analysis uses a kinetic model with a discrete number of conformational states and finds the FRET efficiency value for each state, as well as the state-to-state transition rates that best describe the data. We found that a four-state Markov chain model allowed us to globally account for all experimental datasets (**Figure S8**, see H^2^MM Analysis in methods), with the FRET efficiency values shared among all datasets. This analysis suggested that SR1 samples the same four conformations, which we term ***microstates***, under all conditions.

We used several methods to validate the results of the optimization. First, we performed “photon recoloring” simulations, in which simulated photon trajectories are generated based on the H^2^MM model parameters and the photon arrival times from measured datasets (54) (see H^2^MM Analysis in Methods). The simulated data were then used to generate “recolored” FRET efficiency histograms, which are shown in **Figures 1 B-E** as solid black lines on top of the raw data histograms (green) (see **Figure S6** for recolored histograms of all repeats and conditions). The good overlaps between the experimental and recolored histograms demonstrate that the model parameters found in the H^2^MM analysis describe the smFRET experimental data well. Second, we validated the models using dwell-time analysis (55). In this method, likelihood-weighted dwell-time distributions are generated based on the data and the H^2^MM model parameters (see Weighted Dwell-Time Analysis in Methods), and the mean dwell-time values for the four states are extracted by fitting these distributions to mono-exponential functions. The very good agreement between the dwell-time values obtained from this analysis (**Figure S9**) and those calculated directly from the H^2^MM parameters can be observed in **Table S2** is taken as an additional validation of the analysis results. Further, we performed a segmentation analysis based on the Viterbi algorithm, which generates a realization of the most likely state sequence for each photon trajectory based on the H^2^MM model parameters (see Burst Segmentation in Methods). We picked up only photon trajectories that were assigned to a single microstate and plotted their histograms (**Figure S10)**. The good separation of the microstate histograms validates the H^2^MM assignment. Finally, the presence of submillisecond conformational dynamics was confirmed in a burst-wise fluorescence correlation analysis (see Burst-Wise Fluorescence Correlation Analysis in Methods), showing a characteristic increase of the processed correlation curves at the timescales matching the H^2^MM values (**Figure S11**).

The FRET efficiency values of the four states obtained from the H^2^MM analysis are 0.850 ± 0.002, 0.608 ± 0.005, 0.419 ± 0.006, and 0.213 ± 0.002, and are marked as dashed color-coded lines in **Figures 1 B-E** (FRET values corrected for donor leak and direct excitation of acceptor are ∼0.83, ∼0.55, ∼0.33, and ∼0.09, see H^2^MM analysis in Methods for error estimation). We term the four identified conformational states ***microstates***, labeled numerically according to decreasing order of their FRET efficiency values. The measured and corrected FRET efficiency values of the four microstates can be compared to published conformations of GroEL qualitatively (12, 33, 35). In particular, it is found that microstate 2 matches the structural model of GroEL under the apo condition (**Figure 1 A**, upper), while microstate 3 matches the nucleotide-bound conformation (**Figure 1 A**, middle) and microstate 4 the GroES-bound conformation (**Figure 1 A**, bottom). We did not find an existing structural model that matched microstate 1, probably because it is relatively infrequently populated. The state-to-state transition rates obtained from the H^2^MM analysis are shown in a kinetic scheme in **Figure 2 A-D**. Under all measured conditions, these transition rates correspond to conformational dynamics on the millisecond to submillisecond timescales — three orders of magnitude faster than the bulk ATP hydrolysis rate of SR1 (compare with **Figure S2 A-B**). These fast transition rates provide a quantitative explanation for the emergence of broad FRET efficiency histograms.

**Figure 2:**
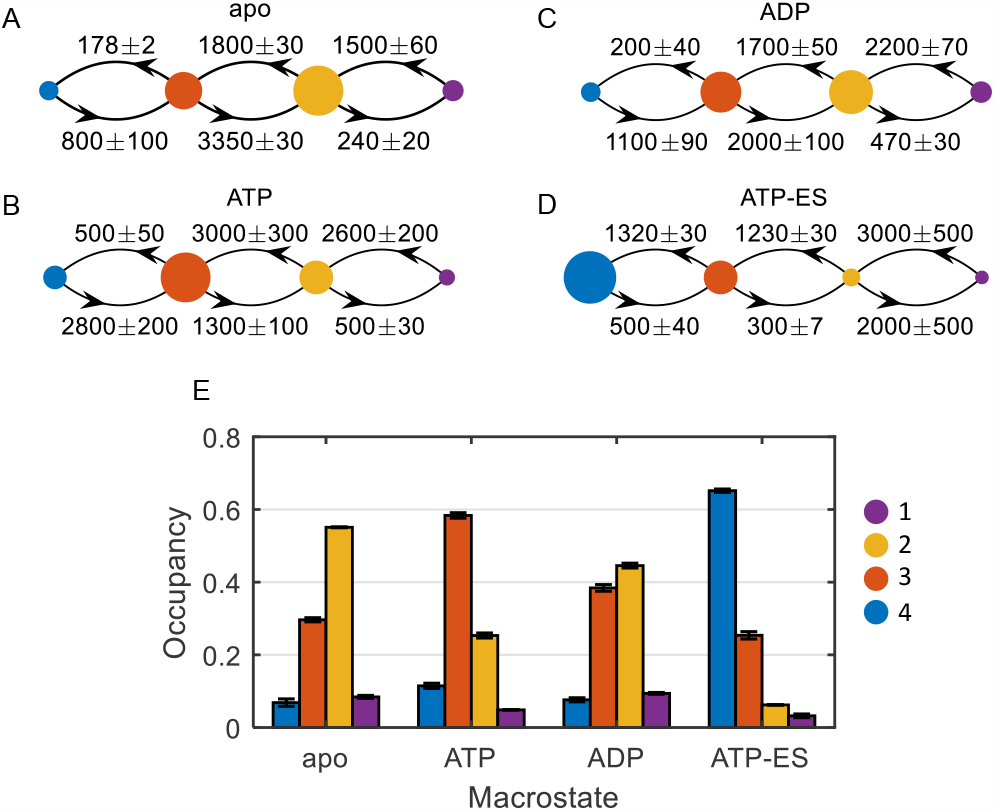
**A-E**) Markov-chain model graphs of various GroEL macrostates. The four microstates are represented with color-coded nodes. The area of each node is scaled to the mean fractional occupancy of each microstate. Transition rates in units of s^-1^ are shown above/below the arrows. **E**) Microstate fractional steady-state populations for each of the macrostates, calculated from the transition-rate matrix. See **Table S3** for numerical values. Errors represent standard error of the mean derived from repeats of the experiment. Microstates are represented with colors as specified.

The steady-state population values for each of the four microstates, directly obtained by diagonalizing the matrix of transition rates, are shown in **Figure 2 E** and **Table S3**. The distribution of microstate probabilities under each condition gives rise to a conformational fingerprint we term a ***macrostate***. In the apo macrostate, the GroEL subunit spends 55.1 ± 0.1% and 29.6 ± 0.6% of the time in microstates 2 and 3, respectively, while microstates 1 (8.4 ± 0.5%) and 4 (7.0 ± 1.0%) are visited less frequently. A notable population shift is seen in the ATP macrostate, resulting in a decrease in the population of microstate 2 to 25.3 ± 0.7% and an increase in that of microstate 3 to 58.4 ± 0.7%. Notice also the slight occupancy increase for microstate 4 (11.5 ± 0.7%). In the ADP macrostate, the observed broad FRET efficiency distribution seen in **Figure 1 D** can be traced to the similar occupancies of microstates 2 (44.6 ± 0.7%) and 3 (38.4 ± 0.9%). In the ATP-ES macrostate, the GroEL subunit resides primarily in microstate 4 (65.2 ± 0.4%) but also in the higher FRET efficiency microstates, with microstate 3 populated 25.4 ± 1.0% of the time.

### Microstate Conformational Dynamics Are Directly Linked to Allosteric Transitions in the GroEL Ring

We investigated the relationship between the observed subunit conformational dynamics and the allosteric transitions of GroEL with respect to ATP by conducting a series of smFRET experiments on samples with increasing ATP concentrations (see smFRET ATP Titration and Burst-Wise Likelihood Analysis in Methods). Since depletion of ATP was expected in experiments with low nucleotide concentrations, the samples contained an enzymatic ATP regeneration system, which ensured that the steady-state conditions were kept throughout the measurements. **Figure 3 A** shows FRET efficiency histograms of SR1 at various ATP concentrations, segmented into the four microstates as discussed above (see also Burst Segmentation in Methods), demonstrating the gradual changes in their populations. To trace the allosteric response, we used a likelihood test to assign each measured molecule to either the apo or ATP macrostate. This procedure was performed for all ATP concentration datasets, and the mean likelihood scores for the apo and ATP macrostates were calculated. Using the scaled score of the ATP macrostate, we generated an ATP saturation curve, as shown in **Figure 3 B**. Remarkably, the plot reproduces the sigmoidal curve familiar from the bulk ATP hydrolysis titration experiment of SR1 (compare with **Figure S2 B**). A fit to a Hill function provides essentially the same Hill coefficient (2.9 ± 0.1) as obtained in the bulk ATP hydrolysis experiment (2.8 ± 0.1).

**Figure 3:**
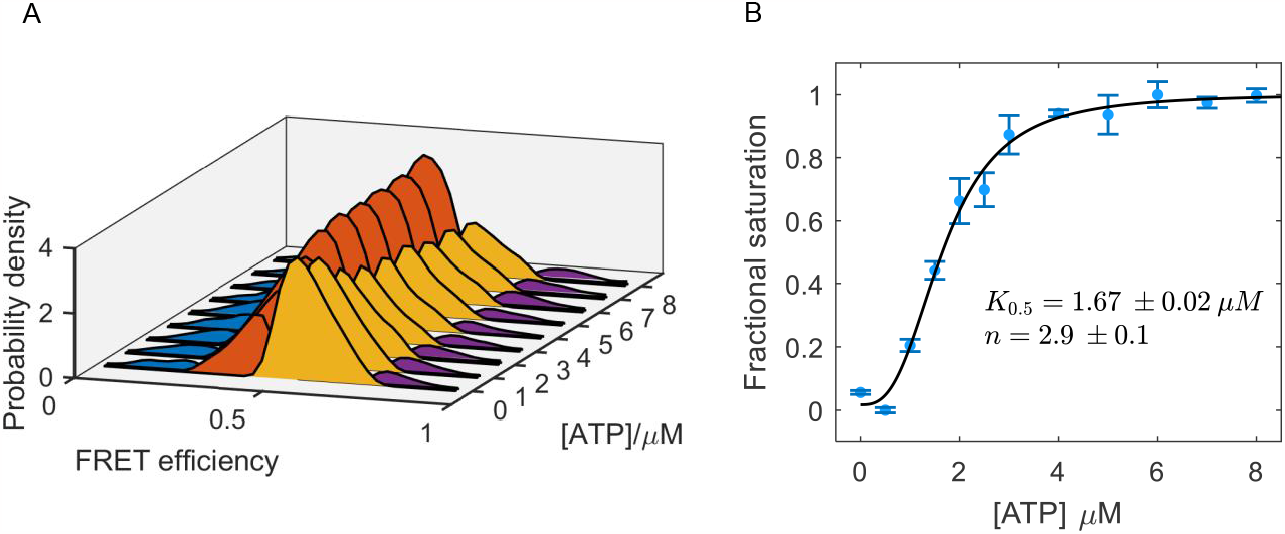
**A**) Segmented histograms of SR1 at different ATP concentrations. The distribution of each microstate is shown with the same color code used in previous figures. Histograms were smoothed using a Gaussian filter with a window size of 5 for visualization purposes. **B**) Fractional saturation vs ATP concertation calculated using the likelihood score. The plot traces a sigmoidal response curve that was fitted to the Hill equation and the resulting parameters are shown in the inset.

## Discussion

In the current study, we directly monitor the conformational dynamics of a functioning GroEL molecule, using time-resolved smFRET spectroscopy to observe fluctuations of a single subunit. Photon-by-photon H^2^MM analysis of the experimental data yields a dynamical model for the conformational transitions. Surprisingly, the GroEL subunit is found to undergo domain motions on the microsecond-millisecond timescale, significantly faster than the functional cycle of the chaperone. These conformational transitions involve four microstates, whose population under each of the probed equilibrium conditions give rise to a dynamical pattern that we term a macrostate. Our modeling approach thus provides a way to characterize the operational phases of the GroEL and its allosteric behavior in terms of the subunit conformational dynamics. Past experiments have probed the kinetics of the GroEL cycle in the bulk (38-41, 49). However, while the conformational dynamics of GroEL have been studied in computer simulations (56-60), a direct, time-resolved measurement of domain motions of the chaperonin has so far been missing.

### Subunit Conformational Dynamics and the GroEL Reaction Cycle

The measurements reported in this study clearly demonstrate that the GroEL subunit adopts multiple conformations under all measured conditions. The characteristic microstate population partition in each macrostate sheds light on the relation between the ensemble of conformations sampled by the GroEL subunit and the functional tasks the machine performs during its reaction cycle. In the non-ligated apo macrostate, the GroEL subunit transitions between the two most populated microstates, 2 and 3, on a time scale of ∼300-500 μs (**Figure 2 A** and **Table S2**). According to the steady-state partition in the apo macrostate (**Figure 2 E Table S3**), the GroEL subunit resides 55% of the time in microstate 2, which, as already mentioned, resembles the subunit conformation seen in the non-ligated structure (**Figure 1 A**). In this closed conformation, the hydrophobic helices of the apical domains face toward the cavity (33), collectively forming a hydrophobic binding surface to which the misfolded protein substrate can bind with higher affinity (18, 20). By dynamically sampling this closed microstate as well as microstate 1, which is likely even more compact, the GroEL subunit is ready for a substrate-binding event. Nevertheless, it still retains the possibility of converting to the more open conformational microstates (3 and 4), which could be essential for subsequent steps such as ATP hydrolysis or GroES binding (61).

Considering the millimolar physiological concentrations of ATP or ADP in the *E. coli* cytosol (62), we can expect the GroEL machine to be mostly bound to these nucleotides, which implies that the machine resides in the ATP and ADP macrostates, respectively. The GroEL subunit continues to exhibit a dynamic behavior in the two nucleotide-bound macrostates, with transitions at ∼300-500 μs (**Figure 2 B-C and Table S2**). Domain motions with similar timescales were reported in simulations probing ligand-induced conformational changes from the apo to the ATP-bound states (58). In the ATP macrostate, as opposed to the apo macrostate, the subunit resides mostly in the microstates with the lower FRET efficiency values (3 and 4), as expected based on structural information (**Figure 1 A**) (34, 35, 63). In such conformations, the apical domain is now extended upwards, and the continuous binding surface, which is prominent in the apo structure, is disrupted. This conformational state diminishes the avidity towards the protein substrate (18), which in turn promotes its subsequent release into the GroEL cavity or back to the solution. Also note that the presence of ATP increases the population of the fully extended, open microstate 4 from ∼7% to ∼11%, which, in line with the conformational selection model (64), could enhance the likelihood of fruitful interaction between the apical domains of GroEL and the loops of the co-chaperonin GroES in subsequent steps of the cycle (12). In the ADP macrostate, which ideally emulates the stage after ATP hydrolysis and the release of phosphate and GroES, the subunit shows similar occupancies for the closed (1 and 2) and open (3 and 4) conformational microstates. Considering the ADP-bound *trans-*ring in the asymmetric reaction cycle of double-ring GroEL (37, 65), such a partition could be advantageous, as it allows the GroEL subunit to readily switch to the closed conformational microstates, allowing it to reform the high-affinity binding surface (20), priming the GroEL ring for a subsequent binding event of a non-native protein substrate.

The behavior of the GroEL subunit in the ATP-ES macrostate is, surprisingly, not static, and pronounced conformational transitions were detected in our experiments, even though the subunit is supposedly “locked” in the long-lived SR1-GroES complex (13, 24, 42, 45) (**Figure S7**). In fact, while the open conformational microstate 4 was predominantly populated in this macrostate, conformational transitions to the more closed microstates were also detected (**Table S2**). A possible explanation for the persistent dynamics in the ATP-ES macrostate is that the apical domain of the subunit occasionally detaches from the GroES loops and is thus free to transition to all other microstates, while the other subunits maintain GroES in place by their collective contributions to avidity.

### The Relation Between Microstate Dynamics and the Allosteric Mechanism of GroEL

The ATP titration curve, generated based on the likelihood score analysis of individual molecules and using the apo and ATP macrostate models, traces the familiar sigmoidal shape for positive cooperativity (**Figure 3 B**). This result directly demonstrates that the conformational motions sampled in the single-molecule experiment of an individual GroEL subunit are associated with the allosteric mechanism of the entire complex. While the functional cycle of GroEL takes place out-of-equilibrium as ATP is continuously hydrolyzed, it is expected that fluctuations in equilibrium will sample the same states on the energy landscape sampled in the cycle. Moreover, the Hill coefficient value obtained in this experiment (2.9 ± 0.1) matches well within error with the value obtained from the bulk ATP titration experiment of SR1 WT (2.8 ± 0.1) (**Figure S2 A**). This striking agreement of the Hill coefficients from two widely different observables – the bulk steady-state ATP hydrolysis rate of the SR1 complex and the local conformational changes of a single labeled subunit– is strong evidence that the allosteric transition in GroEL involves transitions between two states without intermediates, matching the concerted model. An interesting consequence of the local probing of a single GroEL subunit is manifested by the apparent binding constants obtained from fits to the Hill function. In the smFRET ATP titration experiment, the apparent binding constant is ∼1.7 μM, matching the value of the non-labeled 255C/428C SR1 variant but lower than the value of ∼3.5 μM for the WT measured in bulk ATP hydrolysis experiments (**Figure S2 A-B**). This observation can be explained considering that the SR1 construct measured in smFRET experiments consists of one labeled 255C/428C monomer and six WT monomers. Consequently, the apparent binding constant might be determined by the ‘local’ properties of the observed labeled 255C/428C subunit. In contrast, the Hill coefficient – a quantity related to the strength of cooperativity between all the protomers in the complex – is governed by the global conformational behavior of the WT protomer majority.

In line with the findings for the concerted allosteric model, we can treat the dynamical apo and ATP macrostates as representative signatures of the familiar T and R allosteric states of GroEL, respectively. By calculating the interstate equilibrium constants of the two models (readily obtained from the transition rates extracted from the single-molecule experiments), we can generate a closed thermodynamic cycle between the T and R states via the microstates, as shown in **Figure 4 A**. This analysis allows us to estimate which microstate is likely to serve as the preferred gateway for the transition from T to R. It is assumed that transitions from T to R occur within each microstate, i.e., without crossing microstates. The equilibrium constants for the horizontal transitions between the microstates (in the direction from 1 to 4) are shown above and below the arrows, and the equilibrium constants for the vertical T to R transitions are taken relative to the transition from microstate 1, labeled *α*_1_. In comparison to all four possible allosteric transitions, the scheme shows that the transition from the conformational microstate 3, associated with the binding of ATP, is energetically the most favorable. When compared to the transition from microstate 2, the transition from microstate 3 yields an energetic gain of ∼0.8 *kcal* · *mol*^−1^ at the experimental temperature of ∼22 °C. While this contribution is not very high, the thermodynamic cycle demonstrates the importance of subunit conformational dynamics, namely, that a fast exchange between the microstates grants the GroEL subunit more opportunities to reach the optimal microstate for a functional event, in this case, the allosteric transition.

**Figure 3:**
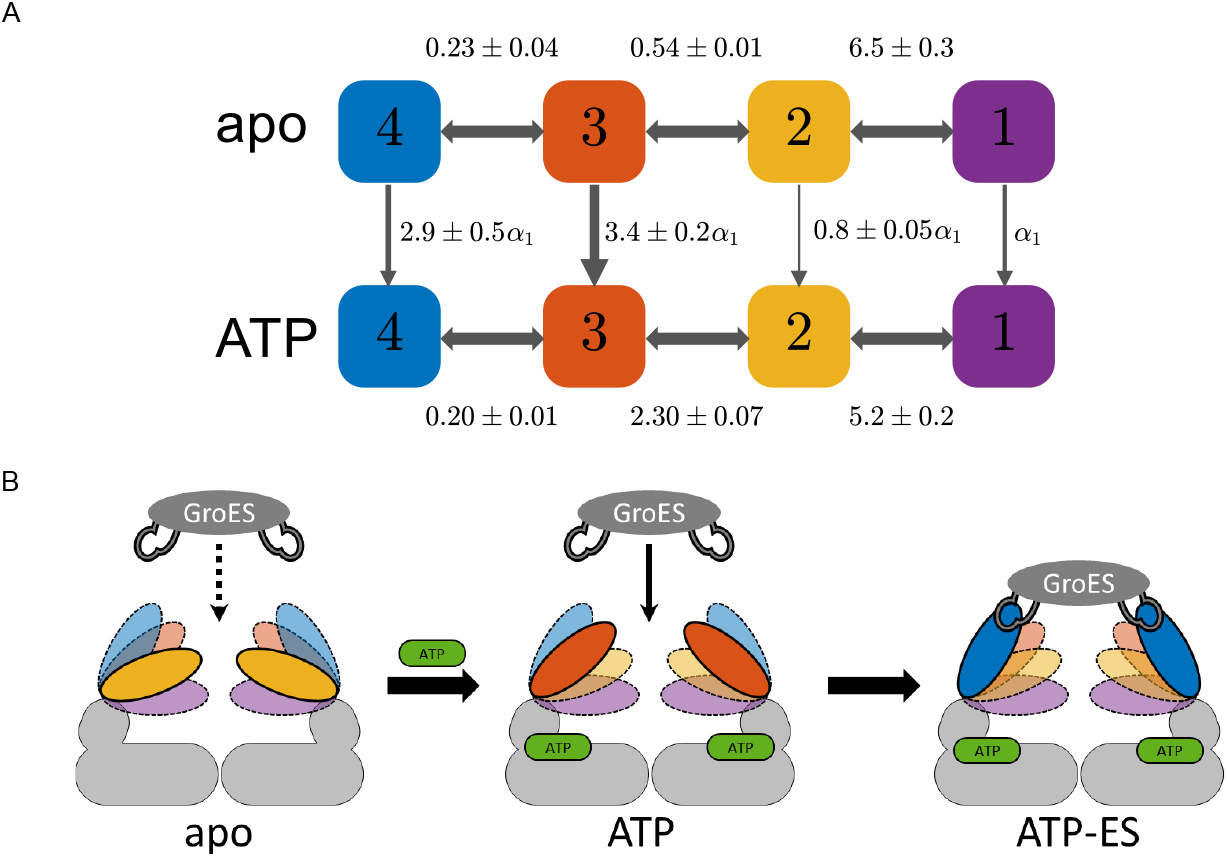
**A**) Closed thermodynamic cycle between the two macrostates apo and ATP. The inter-microstate transitions are shown with horizontal arrows with the adjacent equilibrium constants calculated as *K*_*i*→*i*+1_ = *k*_*i*+1_/*k*_*i*_, where *k*_*i*_, *k*_*i*+1_ are the mean microstate transition rates reported in **Figure 2 A-B**. The apo to ATP transitions are marked with vertical arrows with the corresponding equilibrium constants, which are scaled relative to the transition from microstate 1 marked as *α*_1_ (also represented by arrow width). Mean values are shown with standard errors derived from repeats of the experiment. **B**) Fast conformational motions and conformational selection. Illustrated is the GroEL ring with the equatorial and intermediate domains in gray and apical domain in colors. The four microstates are represented by the different colors and positions of the apical domains. In each macrostate, the most occupied microstate is highlighted with a solid line. In the apo macrostate, the closed microstate 2 is mostly sampled, resulting in low affinity between the apical domains and the GroES mobile loops. In the ATP macrostate, microstate 2 is mostly sampled, together with a slight increase in the occupancy of microstate 1. As a result, the GroES mobile loops can bind with higher affinity to the apical domains, leading to the long-lived ATP-ES macrostate.

### The importance of fast conformational dynamics

As has been shown in the past, fast conformational dynamics have a widespread, fundamental functional role in various biological processes, e.g., in promoting enzyme catalysis (66, 67), molecular machine regulation (68, 69), and protein disaggregation (70, 71). Here we showed that GroEL also exhibits fast conformational motions, shuttling between four microstates, which are associated with various functional aspects in the chaperonin reaction cycle. What could be the significance of fast conformational motions at a submillisecond timescale in the case of GroEL? As a chaperone, GroEL interacts via its apical domains with various non-native protein substrates in a promiscuous manner since unfolded or misfolded proteins typically lack a specific structure (7-10). During the time window of the formation of a collision complex between GroEL and a protein substrate, fast domain motions can facilitate multiple binding and release opportunities between the apical domains and the exposed hydrophobic regions on the protein substrate. This fast conformational motion of the GroEL subunit can also be related to the emergence of asymmetries among the ring subunits, which were directly detected (34) or hinted at (35) in structural studies. As the thermodynamic cycle in **Figure 4 A** implies, fast conformational motions may also be significant for the much slower functional cycle of the chaperone, as the recurrent population of all microstates facilitates, through conformational selection, the transition to the next step in the cycle, e.g., binding of ATP or the co-chaperone GroES. (**Figure 4 B**)

To conclude, the intrinsic ability of the GroEL complex to frequently sample multiple conformational states and modulate its functional capabilities allows it to maintain a continuous capacity to respond to a variety of internal and external molecular events occurring on various timescales in the complex cellular environment. We believe that the mechanistic aspects associated with fast conformational dynamics are a common and fundamental trait in many biomolecular processes, which, with the advent of more sophisticated single-molecule measurement and analysis techniques, can now be readily observed and investigated.

## Materials and Methods

Detailed information on the materials used, protein purification, labeling and reassembly protocols, as well as single-molecule data acquisition and analysis can be found in the provided supporting information section.

## Acknowledgments

We thank Drs. Shay Vimer and Michal Sharon for help with native mass spectrometry and analysis and Dr. Ilia Korobko for providing materials and advice. This work was funded by the European Research Council (ERC) under the European Union’s Horizon 2020 Research and Innovation Programme (grant agreement No 742637). Yoav Barak is the incumbent of Beatrice Barton Research Fellowship.

