## Supplementary Information for "From Microstates to Macrostates in the Conformational Dynamics of GroEL: a Single-Molecule FRET Study"

#### Supporting Information

**This PDF file includes:**

Supporting text

Figures S1 to S11

Tables S1 to S3

SI References

### Supplementary Methods

#### Searching for Labeling Sites Using risFRET

Several factors are to be considered when searching for labeling positions for FRET studies: i) altering the amino acid residues to an orthogonal chemical group for the labeling reaction (in this work, to a cysteine residue) must not disrupt the structural integrity and function of the protein; ii) the labeling positions must be exposed enough to the solution to facilitate the chemical conjugation between the labeling site and functional anchoring group of the dye (in this work, maleimide); and iii) the labeling sites should be selected so that the distance between the donor and acceptor dyes (in this work, AlexaFluor 488 and AlexaFluor 594) is similar to their Förster distance, and changes significantly upon a conformational transition.

To search for labeling-site pairs on a single subunit in the GroEL complex that comply with these considerations, we performed simulations using rotational-isomer model representations of fluorescent labels and their linkers, using our program risFRET, as detailed below. The simulations were based on several double-ring PDB structures of different conformations (1XCK: apo (1), 4AAS: ATP-bound (2), ATP-ES: 1AON (3)). Residues with functional roles were identified based on prior literature information (4) and were excluded from the simulation.

To represent the $C_5$-maleimide fluorescent dyes and their linkers, each dye was modeled in turn on each amino-acid position of a particular PDB model. The dye molecules were represented by a sphere (radius 0.359 nm (5)) attached to the β-carbon of the relevant residue with a $C_{11}$-polyethylene polymer linker. Ten thousand realizations of dye-linker molecules were generated stochastically from the attachment site in a stepwise manner, with the energies of carbon-carbon conformations (anti, gauche+, and gauche, derived from (6)) as Boltzmann statistical weights. Any dye-linker conformations with steric clashes (i.e., with two atoms whose van der Walls radii overlap more than 0.04 nm (7)) were discarded. After generating the dye positions for each site, the expected FRET efficiency value for each pair of residues was calculated as

$$\langle E \rangle = \sum_i \sum_j \frac{p_i q_j}{1 + R_{ij}^6 / R_0^6}, \tag{1}$$

where $R_{ij}$ is the inter-dye distance of each dye-linker realization pair, $i, j$, and $p_i, q_j$, are the Boltzmann statistical weights used in the analysis. A Förster distance, $R_0$, of 0.54 nm was used for the FRET pair AlexaFluor 488 and AlexaFluor 594. The calculated FRET efficiency values were integrated into a score function that weighed together several parameters as follows:

$$score = \Delta E \times (AS_{PDB1}^{res1} AS_{PDB1}^{res2} AS_{PDB2}^{res1} AS_{PDB2}^{res2})^{1/4} \times \{[10 - CS_{res1}][10 - CS_{res1}]\}^{1/2} . \quad (2)$$

$\Delta E$ is the magnitude of the FRET efficiency change between two GroEL PDB models. $AS_{PDB}^{res}$ is a measure of the available space of the linker attached to a labeling site in a specific PDB structure, calculated as the number of non-clashing simulated linker realizations divided by the total number of simulated linkers (10,000 realizations in this analysis). $CS_{res}$ is the evolutionary conservation score for each residue leableing site obtained from the ConSurf webserver (8), which ranges from 1 to 9 from most variable to most conserved. Geometric means of individual volume and conservation scores are calculated to reduce the contribution of extreme values. The higher the final score, the more favorable the labeling site.

Sorting the labeling pairs according to their scores significantly eased the search for appropriate FRET pairs. We initially selected eight labeling-site pair candidates. Preliminary smFRET experiments, conducted as discussed below, demonstrated that the pair E255/D428 showed distinct FRET efficiency changes with respect to ATP and GroES, while the other pairs showed little or no change. We, therefore, adopted this pair for our studies.

#### Purification of GroEL Variants

The following sections describe the purification procedure for the double-ring and single-ring (SR1) GroEL variants. GroES was purified as described in (9). Cysteine mutations on DNA sequences of both double-ring and single-ring GroEL were introduced using standard PCR site-directed mutagenesis. The PCR products were processed using the KLD enzyme mix (M0554S, New England BioLabs) and transformed into DH5-α competent cells. DNA plasmids were extracted from cells using Wizard Plus SV Minipreps (A1465, Promega) DNA purification kit. All purified DNA products were validated by DNA sequencing.

##### GroEL Double-Ring

**Transformation and cell growth:** Plasmids harboring the gene for wild-type (WT) GroEL and cysteine-mutation variants were transformed into TG1 heat-shock competent cells. Individual cell colonies were selected and grown in 0.25 liter of 2xYT growth medium [16 g bacto tryptone (211705, BD Biosciences), 10 g yeast extract (211929, BD Biosciences), 5 g NaCl (0277,

Mallinckrodt Baker) per 1 liter] with 100 µg/ml ampicillin (Formedium, AMP05) at 37 °C under 220 rpm shaking. No induction was required, and the incubation was stopped after ~14 hours.

**Cell lysis:** From this point on, all samples were kept on ice and maintained at 4 °C. The cell suspensions were centrifuged at 6,000 g for 30 min, and cell pellets were resuspended in 20-30 ml cold resuspension buffer [50 mM Tris-HCl (04819620, MP Biomedicals, 10% w/v sucrose (4072, Mallinckrodt Baker), pH 7.5] in new 50 ml tubes. The cell pellets were centrifuged again at 6000 g for 30 min and resuspended in 25 ml lysis buffer [50 mM Tris-HCl, 60 mM KCl (104936, Merck Millipore), 10 mM $MgCl_2$ (M1028, Sigma), 2 mM DTT (11583786001, Sigma), 0.1 mM EDTA (27285, Sigma), pH 7.5], supplemented with 2.5 µl of benzonase (70746, Merck Millipore), and protease inhibitor tablet (04693132001, Roche)). Cells were lysed by sonication (Vibra-Cell VC 650, Sonics) on ice using a microtip at 40% amplitude for 6 min with a 30/60 seconds on/off cycle. The cell lysates were transferred to high-g-tubes and centrifuged at 50,000 g for 1 hour. The protein-containing supernatants were transferred to small beakers.

**Ammonium sulfate precipitation (ASP):** In ASP, GroEL precipitates by salting out from the solution while maintaining its native structure (10). In addition to removing non-precipitating contaminants, the resulting GroEL pellet obtained in ASP can be safely stored at 4 °C for 1-3 days in between purification steps without damaging the protein. The ASP protocol, which was utilized throughout the purification steps, is as follows: 40-45% (w/w) powdered ammonium sulfate (0049, J.T. Baker) is gradually added to the cold protein sample in a glass beaker under light stirring at 4 °C. After 1-2 hours of incubation in the cold room, the sample is transferred to a new tube and centrifuged at 10,000 g for 30-60 min at 4 °C. The supernatant is discarded, and the pellet is kept in the fridge at 4 °C until the next purification step.

**Anion exchange chromatography (AEC) I:** All chromatographic steps described below were performed using an ÄKTA Avant 25 fast protein liquid chromatography (FPLC) system (28930842, Cytiva). The protein pellets from the ASP procedure were resuspended in 40 ml of AEC1 buffer A (50 mM Tris-HCl, 0.1 mM EDTA, 1 mM DTT, pH 7.5) and loaded into a Hi-Prep Q-HP 16/10 column GE28-9365-43, Cytiva) equilibrated with AEC1 buffer A. After loading the sample, a stepwise gradient with AEC1 buffer B (50 mM Tris-HCl, 0.1 mM EDTA, 1 mM DTT, 1 M NaCl, pH 7.5) was applied with gradient steps: 0-24%, 24-40%, and 40-100%. Determination of protein content in collected elution fractions using sodium dodecyl-sulfate polyacrylamide gel electrophoresis (SDS-PAGE) showed that the absorption peak of the second gradient step in the chromatogram contained purified GroEL. GroEL-containing fractions were then collected and processed again by ASP.

**Size exclusion chromatography (SEC):** The protein pellets from ASP were resuspended in ~2.5 ml G10K buffer without $MgCl_2$ (50 mM Tris-HCl, 10 mM KCl, 1 mM DTT, pH 7.5). Buffer exchange was performed on the samples using PD-10 columns (17085101, Cytiva) loaded with the same buffer. The columns used for each GroEL sample were later purged with pure water in order to be reused in the proceeding purification steps. To the eluted samples, $MgCl_2$ and ATP (A2383, Sigma) were added to reach 10 mM and 1 mM concentrations, respectively. This step helps to detach protein contaminants that might still be bound to GroEL. The samples were loaded onto a HiLoad 16/600 Superose 6 prep grade column (29323952, Cytiva), and fractions containing GroEL were collected and processed for ASP.

**Anion exchange chromatography II:** The protein pellets from ASP were resuspended in ~2.5 ml AEC2 buffer A [50 mM MES (M3671, Sigma), 1 mM EDTA, 1 mM DTT, 25% methanol (136805, Bio-Lab), pH 6.0]. Buffer exchange was performed on the samples using PD-10 columns loaded with the same AEC2 buffer A. The samples were loaded onto a monoQ 5/50 GL column (17516601, Cytiva) equilibrated with AEC2 buffer A at 1 ml/min flow. A stepwise gradient with AEC2 buffer B (50 mM MES, 1 mM EDTA, 1 mM DTT, 25% methanol, 1 M NaCl, pH 6.0) was then applied with a gradient step of 0-25%. After contaminant removal, a gradient from 25-100% for 30 min was applied, and eluted fractions containing GroEL (based on SDS-PAGE inspection) were collected and concentrated using a Vivaspin 6 column with a 30 kDa molecular weight cut-off (MWCO) (28932317, Cytiva) to a final volume of ~2.5 ml.

**Acetone precipitation:** Buffer exchange was performed on the samples using PD-10 columns loaded with G10K buffer (50 mM Tris-HCl, 10 mM $MgCl_2$, 10 mM KCl, 1 mM DTT, pH 7.5). The samples were transferred to 30 ml Corex glass tubes (8445, Corex) to prevent the sticking of precipitating pellets. A pre-calculated amount of acetone (1030501, Bio-Lab) was added dropwise to reach 45% (v/v) under light stirring, followed by a ~10 min rigorous stirring using a shaker. The samples were then centrifuged at 30,000 g for 15 min at 4 °C. The supernatant was removed, and the pellets were left to dry at room temperature air for no longer than 5 min. The pellets were then solubilized in a G10K buffer solution, transferred to new tubes, and centrifuged at 20,000 g for 10 min. The supernatants of each sample were transferred to new tubes, and GroEL monomer concentrations were determined using a Cary-300 spectrophotometer (10071600, Agilent Technologies), assuming an extinction coefficient of 10,430 $M^{-1}cm^{-1}$ for the GroEL monomer (11).

All samples were aliquoted and flash-frozen in liquid nitrogen and stored at -80 °C until further use.

**GroEL Single-Ring (SR1)**

**Transformation and cell growth:** SR1 plasmids of double-cysteine and WT variants, containing a poly-histidine tag at the C-terminus, were transformed into *E.coli* BL21 heat-shock competent cells. Individual cell colonies were selected and grown in 0.5 liter Luria broth (LBL0101, Formedium) medium with 100 µg/ml ampicillin at 37 °C under 220 rpm shaking. Upon reaching $OD_{600}$ of 0.6-0.8, induction was initiated with the addition of Isopropyl β-D-1-thiogalactopyranoside (1758-1400, Inalco pharmaceuticals) to 1 mM, and the incubation continued for additional 5-6 hours. From this point, all samples were kept on ice and maintained at 4 °C.

**Cell Lysis:** The cell suspensions were centrifuged at 6,000 g for 30 min, and the pellets were resuspended in 20-30 ml of a cold binding buffer [20 mM Tris-HCl, 500 mM NaCl, 20 mM imidazole (104716, Millipore), 1 mM DTT, pH 8.0], supplemented with $MgCl_2$ to reach 10 mM, 3-4 µl of benzonase, and protease inhibitor cocktail [1 mM PMSF (P7626, Sigma), 0.4 mM benzoamidine (105240250, Arcso Organics), 0.06 mM benzoamide (159330050, Arcos Organics)]. Cells were lysed by sonication on ice, using a microtip at 40% amplitude for 6 min with 30/60 seconds on/off cycle.

**Immobilized metal affinity chromatography:** Cell lysates were centrifuged at 30,000 g for 30 min, and the protein-containing supernatants were loaded onto a 5 ml HisTrap HP nickel column (17524802, Cytiva) pre-equilibrated with the binding buffer. A stepwise gradient with an elution buffer (20 mM Tris-HCl, 500 mM NaCl, 500 mM imidazole, 1 mM DTT, pH 8.0) was applied. Eluted fractions were analyzed with SDS-PAGE on a 15% polyacrylamide gel, and fractions containing pure product were collected and concentrated with a Vivaspin 20, 100 kDa MWCO column (28932360, Cytiva) to a final volume of ~1-2 ml.

**Dimethyl sulfoxide (DMSO) incubation:** To remove protein contaminants, the concentrated samples were supplemented with DMSO (7033, J.T. Baker) to reach a 1:6 ratio and incubated for 2 hours at room temperature under light stirring. The samples were centrifuged at 14,000 g for 10 minutes, and the supernatants were desalted on PD-10 columns equilibrated with G10K buffer (50 mM Tris-HCl, 10 mM $MgCl_2$, 10 mM KCl, 1mM DTT, pH 7.4). Monomer concentrations were determined as above. All samples were aliquoted and flash-frozen in liquid nitrogen and stored at -80 °C until further use.

#### Protein Labeling

This section provides the labeling procedure details for the SR1 E255C/D428C variant. This general protocol was also implemented to label the GroEL double-ring and single-cysteine variants mentioned in this work.

Purified GroEL aliquots were thawed and supplemented with DTT to reach ~5 mM. After 20 minutes of incubation, the samples were desalted into the labeling buffer (50 mM Tris-HCl, pH 7.0) using Bio-spin 6 columns (7326228, Bio-Rad). Alexa Fluor 488 C5-maleimide (donor, A10254, Invitrogen) and Alexa Fluor 594 C5-maleimide (acceptor, A10256, Invitrogen) were added to the labeling reactions to reach a final monomer:donor:acceptor molar ratio of 1:1.2:1.2 with an effective GroEL monomer concentration of ~10 μM. Labeling reactions were performed at room temperature in two successive steps, starting with the addition of the acceptor and one hour of incubation, followed by the addition of the donor. After 2.5-3 hours, the labeling reactions were terminated with the addition of 20 mM DTT followed by 20 minutes of incubation. The samples were then diluted 1:1 with AEC3 buffer A [50 mM MES, 4 M urea (1.08487, Supelco), 1 mM DTT pH 6.0] to disassemble the GroEL oligomers into monomers for the subsequent mixing procedure. The labeled monomer samples were loaded onto a monoQ 5/50 GL column equilibrated with AEC3 buffer A at room temperature. Samples were eluted with a gradient (flow rate: 1 ml/min, 100% in 100 min) of AEC3 buffer B (50 mM MES, 4 M urea, 0.5 M NaCl, pH 6.0) to separate peaks of non-labeled, single labeled, and double-labeled GroEL monomers. The fractions containing double-labeled monomers were identified and collected based on their absorption spectra. The labeling efficiency of the samples was determined spectrophotometrically, and satisfactory dye:subunit ratios of 0.8-1.3 were found for both donor and acceptor dyes. The samples were then aliquoted, flash-frozen in liquid nitrogen, and stored at -80 °C until further use.

#### Reassembly Procedure

To detect FRET signals from only one GroEL subunit in a ring, we prepared GroEL constructs with a single labeled monomer in each complex, using the reassembly procedure shown schematically in **Figure S1**. Our reassembly protocol was based on several published studies (12-19), which utilize the ability of GroEL to disassemble into monomers in the presence of 2-4 M urea and to reassemble back into a functional complex upon removal of urea under reassembly conditions. The idea, first implemented in our lab with the ClpB (20-22), is to reassemble GroEL complexes using a mixture containing labeled monomers and an excess of non-labeled monomers, e.g., in a ratio of 1:100. Assuming that the assembly of GroEL monomers follows a binomial distribution, this ratio would yield a final product, where the fraction of GroEL complexes with only one labeled monomer out of all the labeled complexes is ~94% for the double-ring variant and ~97% for the single-ring variant.

This section details the reassembly procedure for the SR1 E255C/D428C variant. The general protocol was also implemented for the reassembly procedure of the GroEL double-ring and single-cysteine variants mentioned in this work. First, GroEL WT samples in a G10K buffer solution were diluted 1:1 with a urea buffer (50 mM Tris-HCl, 8 M urea, 1 mM DTT, pH 7.4) to reach a final 4 M urea concentration and incubated for 30 min at room temperature. A thawed sample of fully labeled GroEL monomers in 4 M urea in the above buffer was mixed with a GroEL WT monomer sample in a stoichiometric mixing ratio of 1:100. The mixtures were transferred to a GebaFlex dialysis tube of 8 kDa MWCO (D020, Geba) and incubated in 200-300 ml dialysis buffer [50 mM Tris-HCl, 0.6 M ammonium sulfate, 10 mM $MgCl_2$, 2-5 mM ADP (A5285, Sigma), 5 mM DTT, pH 7.4] at room temperature in the dark under light stirring for 10-16 hours. The dialyzed samples were transferred to low-adhesion tubes (1415-2690, USA scientific), centrifuged at 14,000 g for 5 min, and loaded onto a Superdex 200 Increase 10/300 GL column (28990944, Cytiva) equilibrated with G10K buffer. Eluted peaks of assembled GroEL with labeled subunits were separated from non-assembled monomers and other complexes (see **Figure S2 D** for native gel scan of eluted fractions). The final protein and fluorescent dye content of the purified GroEL samples were determined spectrophotometrically, yielding 1-7 μM GroEL oligomers (labeled and non-labeled) and 100-400 nM dye. Samples were then aliquoted, flash-frozen with liquid nitrogen, and stored at -80 °C until further use.

#### Denaturing and Native Gel Electrophoresis

For the inspection of protein samples in denaturing conditions, a 10-15% SDS-PAGE gel was prepared according to the standard protocol with a 200 V voltage and a running time of 50 minutes. For native-gel electrophoresis, native or denatured GroEL samples were loaded onto a 6% polyacrylamide native gel (prepared with the same standard protocol but without adding the denaturing reagent SDS). The samples were loaded onto a gel filled with a native running buffer [25 mM Tris-HCl, 192 mM glycine (07132391, Bio-labs), pH 8.3], with a voltage of 90 V for 2 hours at room temperature. Gel scans of fluorescent labels were performed using a Typhoon FLA 9500 biomolecular imager (GE Healthcare Life Sciences) with 473 nm and 532 nm excitation sources and using 510 nm and 575 nm high-pass emission filters. Gels were stained using standard protocols.

#### Steady-State Kinetic Assays

Steady-state ATPase activity measurements on GroEL constructs were performed using a coupled reaction assay (20, 23) containing pyruvate kinase (PK) and lactate dehydrogenase (LDH), which hydrolyzes nicotinamide adenine dinucleotide (NADH). Samples were prepared with GroEL (100-200 nM oligomers final concentration) in a reaction buffer [50 mM Tris-HCl, 10 mM $MgCl_2$, 10 mM KCl, 0.15 mM NADH (N8129, Sigma), 2 mM phosphoenol pyruvate (PEP, 10108294001, Roche), 1 mM DTT, ~20-30:30-50 units/ml PK:LDH mix (P0294, Sigma), final concentrations] and in the presence of varying ATP concentrations. Triplicate samples were loaded on a 96-well plate, and the NADH absorption at 340 nm was monitored over time with a microplate reader (CLARIOstar, BMG Labtech). After background subtraction, the time-dependent absorption curves were fitted to a linear function. Catalytic rates were calculated by considering the NADH extinction coefficient of 6220 $M^{-1}cm^{-1}$, a cell path length of ~0.59 cm, and the concentration of GroEL in the solution. The generated ATP titration curves were fitted to the Hill equation

$$V(x) = \frac{V_{max}}{1+\left(\frac{K_{0.5}}{x}\right)^n} + b, \qquad (3)$$

where $b$ is a baseline parameter, $V_{max}$ is the maximum reaction rate, $K_{0.5}$ is the apparent dissociation constant, and $n$ is the Hill coefficient. Data analysis and fitting were performed with a Matlab script utilizing a weighted non-linear least-square minimization algorithm (trust-region) and constraining the parameters $n, V_{max}$, and $K_{0.5}$ to positive values.

#### Native-Mass Spectrometry

GroEL samples were treated three times using a Bio-spin 6 column loaded with a buffer containing 150 mM $NH_4OAc$ (238074, Sigma). The samples were then loaded on a gold-coated capillary and introduced into the Q-Exactive-UHMR mass spectrometer for intact mass measurements. The instrument was operated under the following conditions: the inlet capillary temperature was set to 250 °C at a voltage of 1.3 kV, vacuum settings were – fore pump 1.61 mbar, HV 3.23e-9 mbar and UHV 2.35e-10 mbar. Inject flatapole offset and bent flatapole DC were set to 5V and 2V, respectively. No HCD voltage was applied. Measurements were performed at a resolution of 10,000. Raw spectra were converted to MassLynx compatible files by the software Databridge (Waters) and analyzed by the MassLynx program.

#### Fluorescence Anisotropy Measurements

To assess the freedom of motion of protein-attached fluorescent dyes, steady-state and time-resolved fluorescence anisotropy measurements were conducted on single-cysteine SR1 complexes labeled with either the donor or the acceptor. To this end, single-cysteine SR1 variants, 255C and 428C, labeled with Alexa 488 C5-maleiminde (donor) and Alexa 594 C5-maleimide (acceptor), were processed according to the urea reassembly procedure to form heptamers with one labeled subunit per heptamer, as described in previous sections.

**Steady-state measurements:** Single-labeled SR1 samples were diluted in a clean buffer [G10K + 0.01% tween (28320, Thermo Scientific) + 1 mM DTT]. ATP and GroES were added as necessary. The final concentrations in each sample were ~100-140 nM SR1 heptamer (~1-2 nM dye), ~ 1mM ATP, and ~180 nM GroES heptamer when added. Each sample was placed in a clean quartz cuvette (115-F-10-40, Hellma), and steady-state fluorescence anisotropy measurements were performed with a spectrofluorometer (Fluorolog FL3-22, Horiba Scientific) with polarizers in the excitation and emission paths. The excitation/emission wavelengths were set as follows: for the donor-labeled samples, 493/516 nm, and for the acceptor samples, 588/612 nm. The excitation and emission bandpass were adjusted to 5-10 nm. Mean steady-state anisotropy values with G-factor corrections were calculated using the FluorEssence software (Horiba Scientific, see **Table S1**).

**Time-resolved measurements:** The single-labeled SR1 samples were diluted in a clean buffer (G10K + 1 mM DTT) to ~ 1-2 nM concentrations, loaded into a flow cell (see “Single-molecule experiments” for details) and measured on the MicroTime200 microscope as described below, but with the following modifications. Molecules were excited with a polarized laser at either 495 nm or 594 nm and pulsed at a repetition rate of 20 MHz (50 ns) with a power of 20 μW. The emitted photons passed through a polarizing beam splitter cube (Ealing), which split the photons according to their polarization into parallel and perpendicular channels. The split photons passed through emission filters for the donor and acceptor labeled samples, respectively, and their arrival times relative to the laser pulse were recorded. The fluorescence anisotropy decay curve for each sample was calculated using the following equation:

$$R(t) = \frac{I_{\parallel}(t) - I_{\perp}(t)}{I_{\parallel}(t) + 2I_{\perp}(t)}, \qquad (4)$$

where $I_{\parallel}(t)$ and $I_{\perp}(t)$ are the photon arrival time relative to the laser pulse of the parallel and perpendicular components, respectively. The decay curves are shown in **Figure S3**.

#### Preparation of GroEL Complexes with the ATP Analog BeFx

**Symmetric double-ring football complex**: A sample of labeled double-ring GroEL 255C/428C bound to GroES on both rings, termed a “football complex”, was prepared by adding the following ingredients in this order: 48 μl pure water, 25 μl 4X folding mix (200 mM Tris-HCl, 400 mM KCl, 200 mM $MgCl_2$, pH 7.5), 0.1 μm filtered and fresh 1 M DTT, 15 μl labeled double-ring GroEL construct (~400 nM oligomer, ~4 nM labeled subunits) and 1.2 μl GroES (~163 nM oligomer). After 15 minutes of incubation at room temperature, 10 μl of X10 activation mix was added [550 mM $Na_2SO_4$ (71962, Fluka analytical), 110 mM NaF (106449, Merck), 10 mM $BeSO_4$ (14270, Sigma)]. The sample was mixed, and 1 μl of 100 mM ATP was added. The final concentrations were: double-ring GroEL ~60 nM, GroES ~150 nM, and 1 mM ATP. The structural integrity of the prepared, labeled GroEL football complex was confirmed on a 6% native gel.

**SR1 samples:** Labeled SR1 255C/428C samples bound to ADP-BeFx were prepared using the same procedure as above, but with the addition of ADP instead of ATP. Samples with or without GroES were prepared.

For smFRET experiments, all prepared samples were diluted to picomolar concentrations in the same prepared buffer and measured as described in the following sections.

#### Single-Molecule Experiments

**Flow cell preparation:** Flow cells for single-molecule experiments were prepared by first etching glass slides (Precision cover glasses thickness No. 1.5H, 24x50 $mm^2$, 0107222, Marienfeld) and coverslips (Cover glasses square, 18X18 $mm^2$, 0101030, Marienfeld) in fresh 10% hydrofluoric acid solution (100338, Supelco) in a sonication bath for 40 seconds. The slides and coverslips were then rinsed with copious amounts of pure water and dried with a gentle nitrogen stream. To form a flow cell with a gap, pairs of slides and coverslips were placed together, separated by two thin parafilm strips (HS234525D, Heathrow Scientific). The flow cells were placed in a sealed glass container with a cell holder and incubated at 115 °C for 15-40 min, causing the parafilm to adhere to the glass and seal the rims of the flow cell.

**Unilamellar vesicles**: A suspension of unilamellar vesicles made of chicken egg L-α-phosphatidylcholine (egg-PC) was used to form a supported lipid bilayer on the glass surfaces in order to passivate them and reduce protein binding. The vesicle suspension was prepared by first solubilizing egg-PC powder (840051P, Avanti) in tert-butanol (471712, Sigma) to 20 mg/ml end concentration in a clean glass tube. The lipid suspensions were aliquoted into low-adhesion tubes, sealed with punctured caps, and flash-frozen with liquid nitrogen. The samples were then placed in a lyophilizer for 4-16 hours to remove the tert-butanol by sublimation. The tubes containing dry egg-PC films were then sealed with caps and stored at -20 °C until further use. A sample of unilamellar vesicles was prepared by hydrating a lipid film sample with 450 μl of G10K buffer and extruding the suspension through a 0.1 μm AnoTop filter (6809-112, Cytiva) attached to two syringes back and forth 37 times at room temperature. The prepared unilamellar lipid samples were kept at 4 °C and used for no longer than two weeks.

**Residual contaminants removal from buffers and reagents:** All buffers used in the single-molecule experiments were treated with active charcoal (242241, Sigma) for 12-16 hours and later filtered with 0.2 μm filter (SLGVV255F, Millipore) to remove any fluorescent contaminants. DTT, ATP, and ADP stock solution were prepared with treated buffers, filtered with a 0.1 μm filter, aliquoted to small volumes, and stored at -20 °C or -80 °C.

**Loading samples into flow cell:** For each measurement, a flow cell was first loaded with 40 μl of G10K buffer, followed by the addition of 40 μl of vesicle suspension and a 5 minutes incubation. The cell was then washed with 800 μl of G10K + 1 mM DTT buffer. GroEL samples were prepared by thawing a fresh 2-3 μl aliquot of labeled GroEL and performing two successive dilutions into a final 100 μl G10K + 1 mM DTT buffer to a concentration of 50-100 pM of labeled molecules. Additional substrates like ADP, ATP, or GroES were added to the final sample when necessary. Finally, the passivated flow cell was loaded with 60-100 μl of the prepared sample, sealed with silicone grease (107746, Supelco) to prevent sample evaporation, and loaded on the microscope.

For the smFRET ATP titration experiments, samples were prepared as follows: Stocks of ATP in G10K buffer were diluted with a 10X regeneration buffer containing PK (P1506, Sigma) and PEP (final G10K regeneration buffer: 2 mM PEP, 20-100 U/ml PK, 1mM DTT, pH ~7.0). The labeled SR1 was added in the last step to reach pM concentrations.

**Optical Setup**: Single-molecule measurements of freely diffusing molecules were conducted on a MicroTime200 microscope (PicoQuant) using a pulsed-interleaved excitation (PIE) scheme (**Figure S4**). Laser pulse sequences at two wavelengths, 485 nm for the donor (LDH-D-C-485, PicoQuant, 50 μW) and 595 nm for the acceptor (LDH-D-TA-595B, PicoQuant, 10 μW) for the acceptor, were driven and synchronized by a Sepia PDL 828 module (PicoQuant) at a repetition rate of 20 MHz (50 nanoseconds pulse sequence cycle). The two lasers were coupled to an optical fiber, collimated, and reflected through a 488/595nm excitation dichroic mirror (Ex-DM, zt473/594rpc, Chroma) into a 60X water objective (UPlanSApo Superacpochromat, Olympus), which focused them ~10 μm into the solution of the measured sample. Emitted fluorescence photons passed through the same objective and the excitation Ex-DM, focused to a 50 μm pinhole and recollimated into an emission dichroic mirror (Em-DM, FF580-FDi01, Semrock) to separate donor and acceptor photons into two channels. Photons passed through emission filters (F-D for the donor channel, 520/35 nm, BrightLine, Semrock; F-A for the acceptor channel, ET-645/75 nm, Chroma) and focused into single-photon avalanche photodiodes (SPAD, SPCM-AQR-14-TR, Excelitas), connected to HydraHarp400 photon counting module, operating at T3 mode. The arrival times of photons relative to the pulses generating them and relative to the overall duration of the experiment were collected and stored using the software SymPhoTime (PicoQuant).

#### Data Analysis

All data analysis procedures were conducted using home-built MATLAB scripts, which can be provided upon request.

**Burst identification and filtration:** To facilitate the identification of photon bursts in a smFRET experiment, characterized by a high photon flux as labeled molecules diffuse through the confocal volume of the microscope, the difference in arrival times between each pair of successive photons in the data, Δt, was first smoothed with a moving average mask of the size of 15 photons to reduce the noise in the search process. This procedure effectively allowed us to analyze the local photon flux at each recorded time point in the experiment. A burst was defined as a stream of successive photons, all having Δt values below 50 μs, with at least 30 consecutive photons with Δt values below 10 μs. The raw FRET efficiency and stoichiometry of each burst were calculated using the photon arrival times relative to the sync-pulse, allowing the separation of double-labeled molecules from donor-only and acceptor-only molecules, as follows:

$$E = \frac{N_A^D}{N_D^D + N_A^D}, \quad (5)$$

$$S = \frac{N_D^D + N_A^D}{N_D^D + N_A^D + N_A^A}, \quad (6)$$

where $N_D^D$ and $N_A^D$ are the number of photons detected in the donor and acceptor channels, respectively, after the donor excitation pulse and photon $N_A^A$ is the number of photons detected in the acceptor channel after the acceptor excitation pulse. Corrected FRET efficiency histograms were generated following this reference (24). Background corrections were performed per burst by subtracting the product between the mean background rates of the experiment with the burst duration (typical values ~0.5-1 photons/millisecond per channel). Corrections for donor photon leakage into the acceptor channel and direct excitation of the acceptor by the donor laser were performed using the mean FRET efficiency of the donor-only population and the mean stoichiometry of the acceptor-only population, respectively. A correction ($\gamma$) factor was estimated by fitting a 2D-PIE histogram from a measurement showing multiple FRET efficiency populations to the stoichiometry-FRET relation shown in (24). For this fit, we used a dataset of SR1 in the presence of 1 mM ADP and GroES (see **Figure S7 B**, blue) and obtained a $\gamma$ factor of ~1. After applying these corrections, photon bursts of molecules labeled with both donor and acceptor dyes and containing a minimum of 50 photons (originating from the donor excitation pulse) were selected for further analysis and the generation of FRET efficiency histograms.

Prior to $H^2MM$ analysis, the selected bursts of each dataset were subject to a filtering process to remove time regions in the photon stream where acceptor blinking or dye photobleaching occurred by inspecting the unsmoothed inter-photon times, Δt, of photons originating from the donor and the acceptor pulses separately. For each burst, time segments with either donor or acceptor photons with a Δt value above 50 μs were split, creating new photon trajectories with new FRET efficiency and stoichiometry values. After filtration, 6000-10000 bursts with ~0.5 stoichiometry with a minimum of 30 photons were collected per dataset.

**$H^2MM$ analysis:** Each set of bursts was subjected to $H^2MM$ analysis with 5-10 initial guesses as described in (25) This analysis is based on a photon-by-photon likelihood maximization algorithm, which yields three sets of parameters: i) the initial state probability of each trajectory, ii) the probability for emitting a particular signal per state, in our case, a donor or an acceptor photon, and iii) a state-to-state transition probability matrix. The physical parameters from the measured system are extracted as follows. First, the FRET efficiency value for each state are directly obtained from the probability of emitting acceptor photons in that state. The FRET efficiency values, $E$ , obtained in this manner are corrected using the following relation:

$$E_{corr} = \frac{E - (1-E)l - d\,\frac{1-s}{s}}{1 - d\,\frac{1-s}{s}}, \qquad (7)$$

where $l$ is the leak factor, $d$ is the direct excitation factor, and $s$ is the mean stoichiometry of the measured bursts, as obtained from the PIE analysis described in previous section. Second, to calculate the state-to-state kinetic transition rates, $k_{ij}$, we use the approximation $k_{ij} \cong \frac{p_{ij}}{\Delta t}$, where $p_{ij}$ is the transition probability obtained in the H$^2$MM analysis. This approximation is based on the fact that the time step $\Delta t$ fulfills the condition $\Delta t \ll \frac{1}{k_{ij}}$. Finally, the occupancy of each microstate at equilibrium is obtained from the eigenvector of the transition probability matrix that corresponds to an eigenvalue of 1 (26).

**Histogram "recoloring":** To check if the model parameters obtained from the H$^2$MM analysis successfully describe the photon data, we perform a photon "recoloring" procedure as introduced in (27). In this procedure, the photons within each photon trajectory (burst) measured in the experiment are "stripped" of their color, i.e., the donor or acceptor channel assignment, and only the photon arrival times are kept. Then, a Markov chain state sequence is simulated on the photon arrival times using the parameters obtained in the H$^2$MM analysis, effectively assigning new colors to each photon. FRET efficiency histograms of the recolored simulated photon trajectories are generated and compared to the real data histogram. A good overlap indicates that the converged H$^2$MM parameters describe the data well.

**Model selection:** In the framework of classical HMM analysis, several statistical criteria can be used to determine the minimal number of states to fit a data set, such as the Bayesian information criterion (BIC) (28) and the Akaike information criterion (AIC) (29). However, when using such measures on photon data analyzed in the H$^2$MM analysis, these statistical scores do not converge to a definitive answer (30, 31). We therefore utilized empirical tests for model selection. Preliminary H$^2$MM analysis was performed on datasets using models with 2 to 6 states with either full inter-state connectivity or using a chain architecture. We estimated the suitability of the tested models by comparing the root mean square difference (RMSD) between the FRET efficiency histogram values of the real data and of recolored realizations (**Figure S8 A**). These scores improved in general as the number of states increased, but models with 4 states and above resulted in the appearance of states with the same FRET efficiency value, a sign of redundancy of the model (**Figure S8 B**). Using a fully connected model resulted in low transition rates between non-neighboring states, pointing towards a chain architecture. These tests indicated that a four-state chain model is a suitable Markov model that best describes the collected data.

**Global analysis:** In the case of global H$^2$MM analysis, the FRET efficiency values of the microstates were shared between several analyzed datasets. This was achieved by calculating the re-estimated parameters for the emission probabilities by sharing the auxiliary γ-parameter

(35) across all datasets, while the other parameters, the prior vector and the transition probability matrix, were updated for each dataset individually. In the global $H^2MM$ analysis, we used 2-3 data sets independently measured for each condition (apo, ATP, ADP: 3 repeats, ATP+ES 2 repeats). The parameters for transition rates and equilibrium populations were not shared between data sets, and we report mean and standard error values calculated from these. In order to estimate the errors in the FRET efficiency values, which were globally shared between data sets, we performed additional analysis runs on two groups, each containing apo, ADP, ATP and ATP+ES datasets.

##### Weighted Dwell Time Analysis

This analysis is based on (32). Consider a measured photon trajectory and the parameter set obtained from the $H^2MM$ analysis. The "weighted dwell time" algorithm collects the states' dwell times for ***all*** the possible Markov state realizations in the photon trajectory and weights each realization by the corresponding likelihood values, calculated using the $H^2MM$ model parameters. This process is carried out for every photon trajectory in the dataset to generate likelihood-weighted dwell time distributions for each Markov state. The dwell-time distributions are then corrected using the Kaplan-Meier estimator (33) to censor state segments that terminate due to the end of photon trajectories rather than due to a jump to a different state. Next, we fit the cumulative sum of the dwell-time distribution of each state to a mono-exponential function $f = (1 - e^{-t/\tau})$ and extract the mean dwell-time values, which can be compared to those calculated directly from the $H^2MM$ parameters (see **Figure S9** and **Table S2**).

##### Burst Segmentation

As one way of validating that the $H^2MM$ model correctly traces the dynamics in photon bursts, a photon trajectory segmentation analysis was performed using the HMM Viterbi algorithm (32). This algorithm generates the most likely state trajectory for each photon trajectory, based on the model parameters obtained from the analysis. Based on the microstate assignments generated by the Viterbi algorithm, segmented FRET efficiency histograms of photon trajectories with no state-to-state transitions were generated and plotted (**Figure S10**). The segmented histograms of each state were normalized to match the raw data histogram.

##### Burst-Sise Fluorescence Correlation Analysis

We performed a burst-wise photon correlation analysis to identify the presence of fast GroEL subunit conformational dynamics on photon dataset of SR1 under the four measured conditions.

To obtain information on longer timescales, we performed the burst identification procedure as described above (see Burst identification and filtration section) by setting a Δt threshold of 50 μs but without demanding a Δt threshold of 10 μs. A correlation analysis was performed with a home-built code utilizing the xcorr MATLAB function, based on ideas presented in (34). For a given smFRET dataset containing bursts of molecules labeled with both donor and acceptor dyes, the correlation functions between photons of channels $i$ and $j$ for a time shift $\tau$, $G_{ij}(\tau)$, were calculated with the following equation:

$$G_{ij}(\tau) = \frac{\frac{\sum_{b^*} \sum_{t=0}^{T_{b^*}-\tau} I_{b^*}^{i}(t) \cdot I_{b^*}^{j}(\tau+t)}{\sum_{b^*} T_{b^*}-\tau}}{\frac{\sum_b N_b^i \sum_b N_b^j}{\left(\sum_b T_b\right)^2}}, b^* \in \{b | T_b - \tau > 0\}. \quad (8)$$

The subscript $b$ denotes a burst in the dataset and $T_b$ is the burst length in arbitrary bin time units (in our experiments, a time bin is 50 nanoseconds). The subset of bursts denoted as $b^*$, corresponds to bursts whose lengths are longer than $\tau$. This distinction deals with the variability in the burst lengths in the dataset. $I_b^i(t)$ is a time-binned photon time trace of channel $i$ of burst $b$, which has the value of 1 if a photon is in time bin $t$ and 0 otherwise. $N_b^i$ is the total number of photons in channel $i$ of burst $b$. The term in the numerator essentially calculates the total number of intersecting photon pairs in all the bursts for the time shift $\tau$, averaged over the total sampling time bins of all bursts considered. The term in the denominator is the normalization according to the mean intensity of each channel.

We calculated for each dataset two correlations functions: $G_{DD}(\tau)$ the auto-correlation function of the donor channel and $G_{DA}(\tau)$, the cross-correlation between the donor and acceptor channels. Both terms contain information on intensity fluctuations due to the diffusion of molecules during a burst and due to conformational dynamics. By calculating the ratio between the two correlation functions, $\frac{G_{DA}(\tau)}{G_{DD}(\tau)}$, the signal from diffusion could be eliminated, resulting in a clearer representation of the signal from conformational dynamics and photophysical effects such as triplet dynamics. The ratios between the correlation functions were smoothed using a logarithmic mean moving window (each window lag time is increased by a factor of 2 compared to the previous window) and are shown in **Figure S11.**

#### Burst-Wise Likelihood Analysis to Assign the Macrostates of Bursts

We performed smFRET experiments with increasing ATP concentrations in a buffer containing an ATP regeneration system (see previous Sample preparation section). For each ATP concentration, 2-3 measurements were conducted. In the burst-wise likelihood analysis, two likelihood values were calculated for each recorded photon burst, based on the parameters of the HMM models corresponding to the apo and ATP macrostates. This calculation allowed us to quantify to what extent a detected burst resembled each of these macrostates. The two likelihood values were calculated using the $H^2MM$-modified forward algorithm (25) and were normalized to their sums, yielding a final score with a value between 0 and 1. The mean likelihood scores based on all bursts in each dataset were then calculated. The likelihood score for the ATP macrostate as a function of ATP concentration is plotted in Figure 3 and fitted to the Hill function (eq. 3).

#### Labeling Controls on the Native Cysteine Residues of GroEL

To validate that the native cysteines of GroEL have a low labeling rate, as previously reported (35-37), we conducted control smFRET experiments on three SR1 constructs. SR1 WT and the single-cysteine SR1 variants, 255C and 428C, were labeled with Alexa 488 C5-maleiminde (donor) and Alexa 594 C5-maleimide (acceptor) as described in the previous sections. The monomer:donor:acceptor ratio for this control experiment was 1:0.5:0.5, with an effective GroEL monomer concentration of ~10 μM. Non-reacted dyes were removed using Bio-spin 6 desalting columns using a 50 mM Tris-HCl (pH 7.4) buffer. The labeled SR1 monomers were then subject to the same urea reassembly procedure described above, mixing labeled GroEL monomers with GroEL WT in a 1:100 stoichiometric ratio. The purified assembled SR1 complexes were flash-frozen in liquid nitrogen and stored at -80 °C until further use. smFRET experiments were performed on the three reassembled labeled SR1 constructs: WT, 255C, and 428C, as described in the following sections, and normalized histograms of corrected stoichiometry were generated (see **Figure S2 F**).

### Supplementary Figures

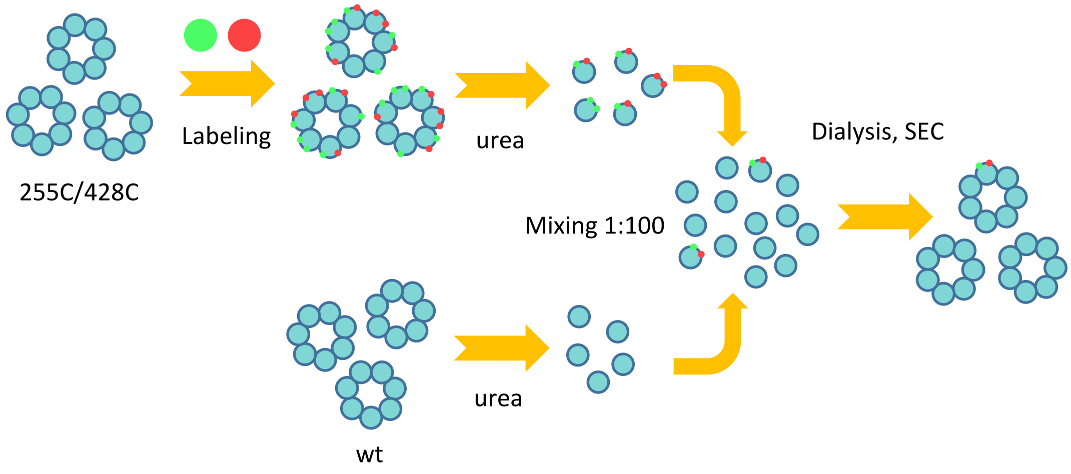


**Figure S1**: Preparation of GroEL constructs by the urea-reassembly procedure. GroEL or SR1 (cyan) with double-cysteine mutations 255C/428C is labeled with donor and acceptor dyes (green and red). The labeled GroEL subunits are separated and mixed with non-labeled WT GroEL subunits in the presence of 4 M urea in a 1:100 ratio. Urea is removed, and GroEL monomers assemble into complexes by dialysis in a reassembly buffer. After an additional SEC step, GroEL complexes containing a single labeled subunit were obtained. A mixing ratio of 1:100 guaranteed that ~97% of the final labeled product molecules consisted of complexes with only a single subunit labeled with fluorescent dyes.

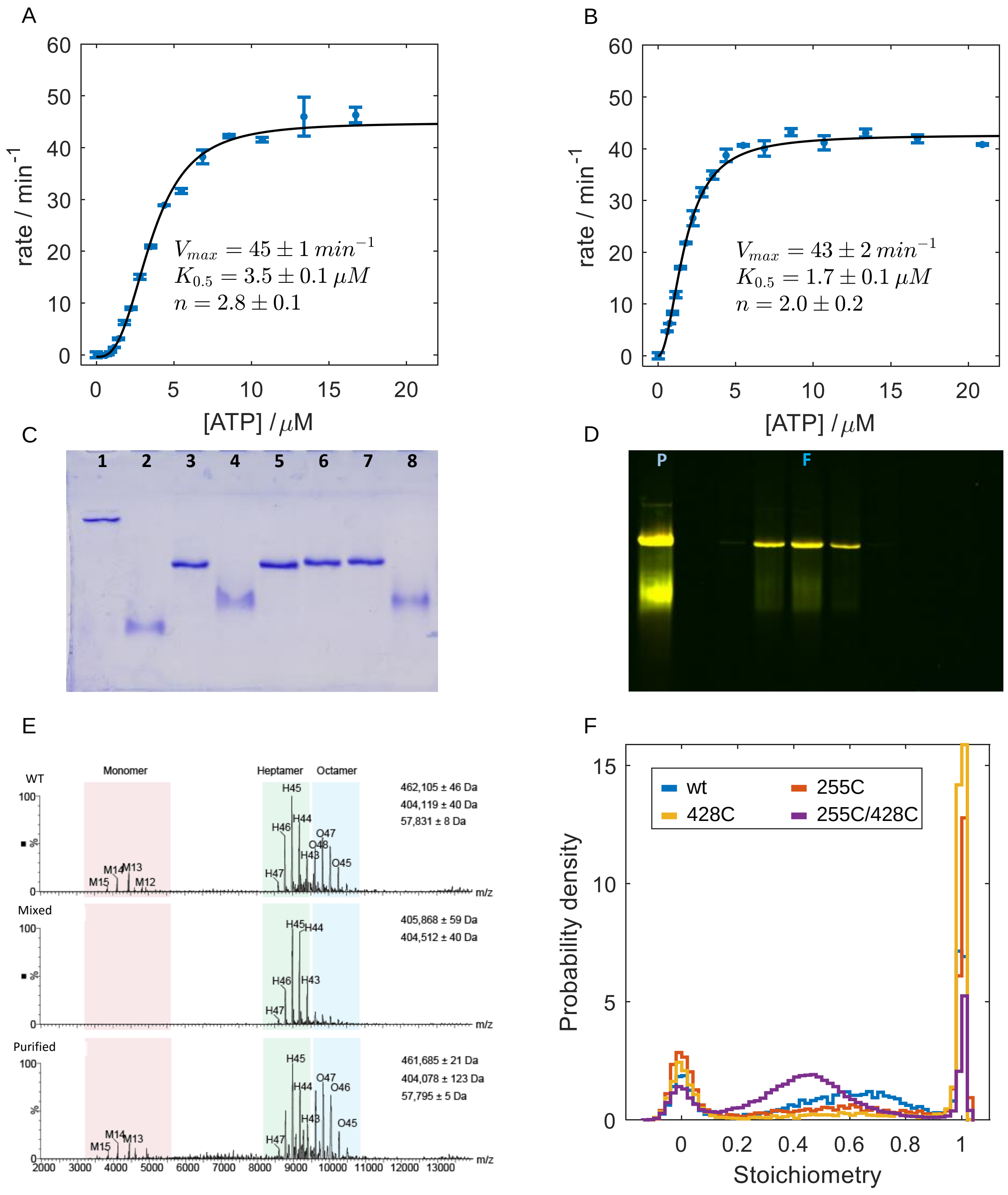


**Figure S2**: Control experiments on SR1 variants. **A-B**) Bulk ATPase activity of SR1 WT (**A**) and SR1 E255C/D428C (**B**). Plots of ATP hydrolysis rate as a function of ATP concertation were fitted to the Hill equation eq.3. The parameters are shown with standard errors based on triplicates. **A-B**) Validation of structural integrity of GroEL complexes. (**C**) 6% native gel scan showing the structural integrity of complexes of different GroEL variants. Lane 1: double-ring WT. Lane 2: double-ring WT monomers incubated in 4 M urea. Lane 3: SR1 WT. Lane 4: SR1 WT monomers incubated in 4 M urea. Lane 5:

SR1 255C. Lane 6 SR1 428C. Lane 7: SR1 255C/428C. Lane 8: SR1 255C/428C monomers incubated in 4 M urea. The similar migration rates of the SR1 WT and cysteine variants confirm the integrity of the heptameric complex. Variations in migration patterns of particles with similar molecular weight can be attributed to charge differences due to the introduced mutations. (**D**) 6% native gel scans of the double-labeled 255C/428C SR1 variant after the urea reassembly procedure. Lane P: product after dialysis and before separation, showing the presence of aggregates, assembled complexes, and monomers. Lanes F: collected fractions of heptameric SR1 particles after size exclusion gel filtration. (**E**) Native mass spectra of SR1 samples. Top panel: SR1 WT. Middle panel: labeled SR1 255C/428C construct after the urea reassembly procedure. Bottom panel: purified 255C/428C (non-labeled). The purified SR1 samples contain, in addition to assembled heptamers (~400 kDa), traces of monomers (~57 kDa) and octamers (~460 kDa). In the reassembly procedure, SR1 monomers assemble into heptamers, and unwanted specimens are removed. **F**) Labeling controls on GroEL native cysteine residues. Normalized corrected stoichiometry histograms of labeled SR1 variants, which were labeled under the same conditions and subject to the reassembly procedure. Bursts with stoichiometry between 0.18-0.9 are assigned to SR1 complexes labeled with both a donor and an acceptor dye. The labeled WT variant (blue) shows a population (~50%) of double-labeled molecules due to non-specific native cysteine labeling. However, when a single mutant cysteine, either 255C (red) or 428C (yellow), is introduced to SR1, the double-labeled population decreases to ~27% and 12%, respectively. This result indicates that the inserted cysteine residues are more reactive than the native cysteines, thereby increasing the population single-labeled molecules. For comparison, the histogram of the double-labeled variant 255C/428C under apo conditions is shown (purple). Thus, we can conclude that labeling the double-cysteine variant under the same conditions as above yields a negligible fraction of SR1 molecules with labeled native cysteine and that the collected FRET signals in our experiments originate from GroEL subunits labeled at positions 255C/428C.

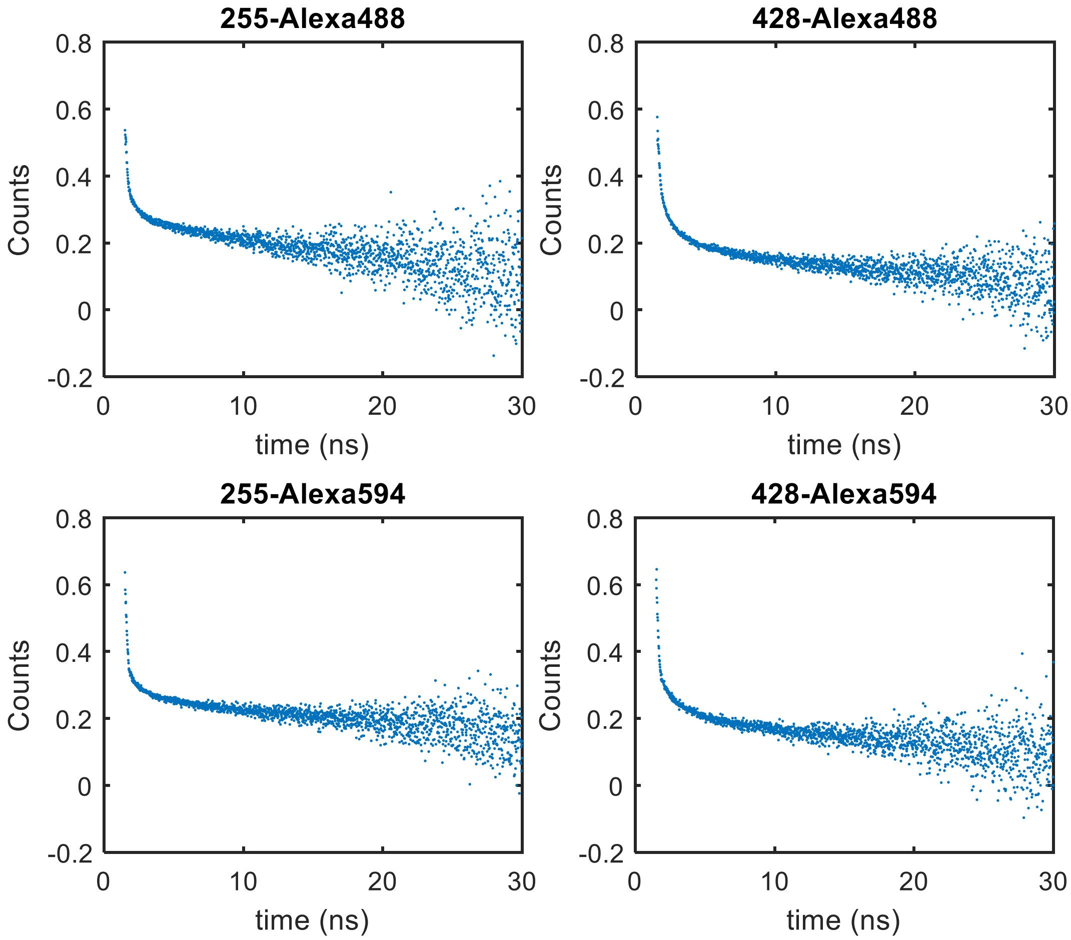


**Figure S3:** Time-resolved anisotropy decays of single-cysteine SR1 construct labeled with the donor or acceptor. The anisotropy decays were calculated according to equation (3). The fast decay component indicates that the dyes at the labeling positions are not significantly constrained, while the slower decay is attributed to the rotational motion of the protein complex.

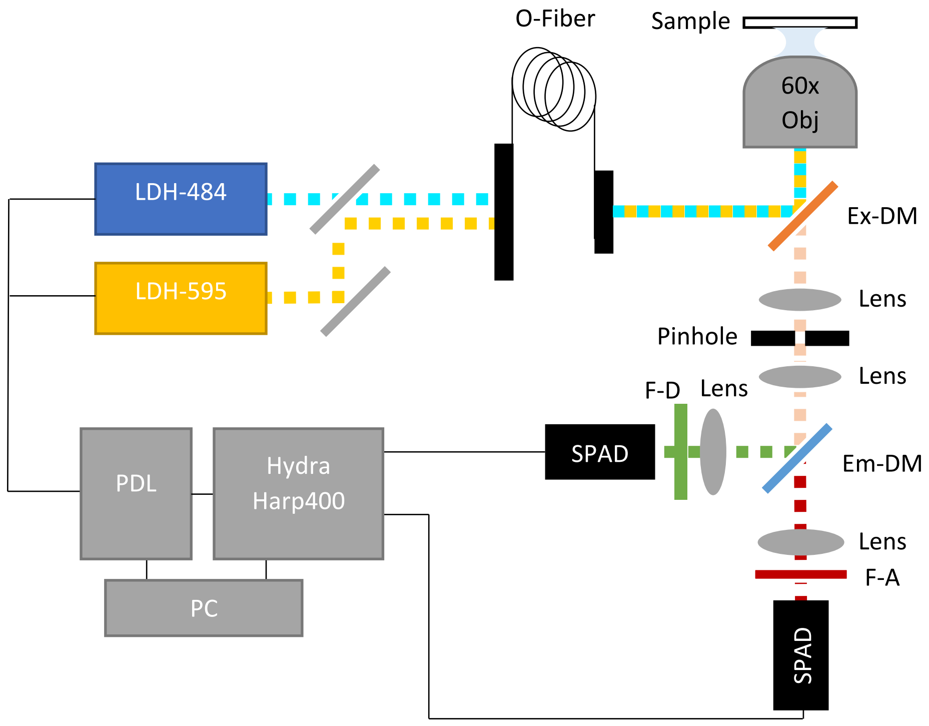


**Figure S4:** Scheme of the MicroTime200 optical setup. The components are abbreviated as described in the text.

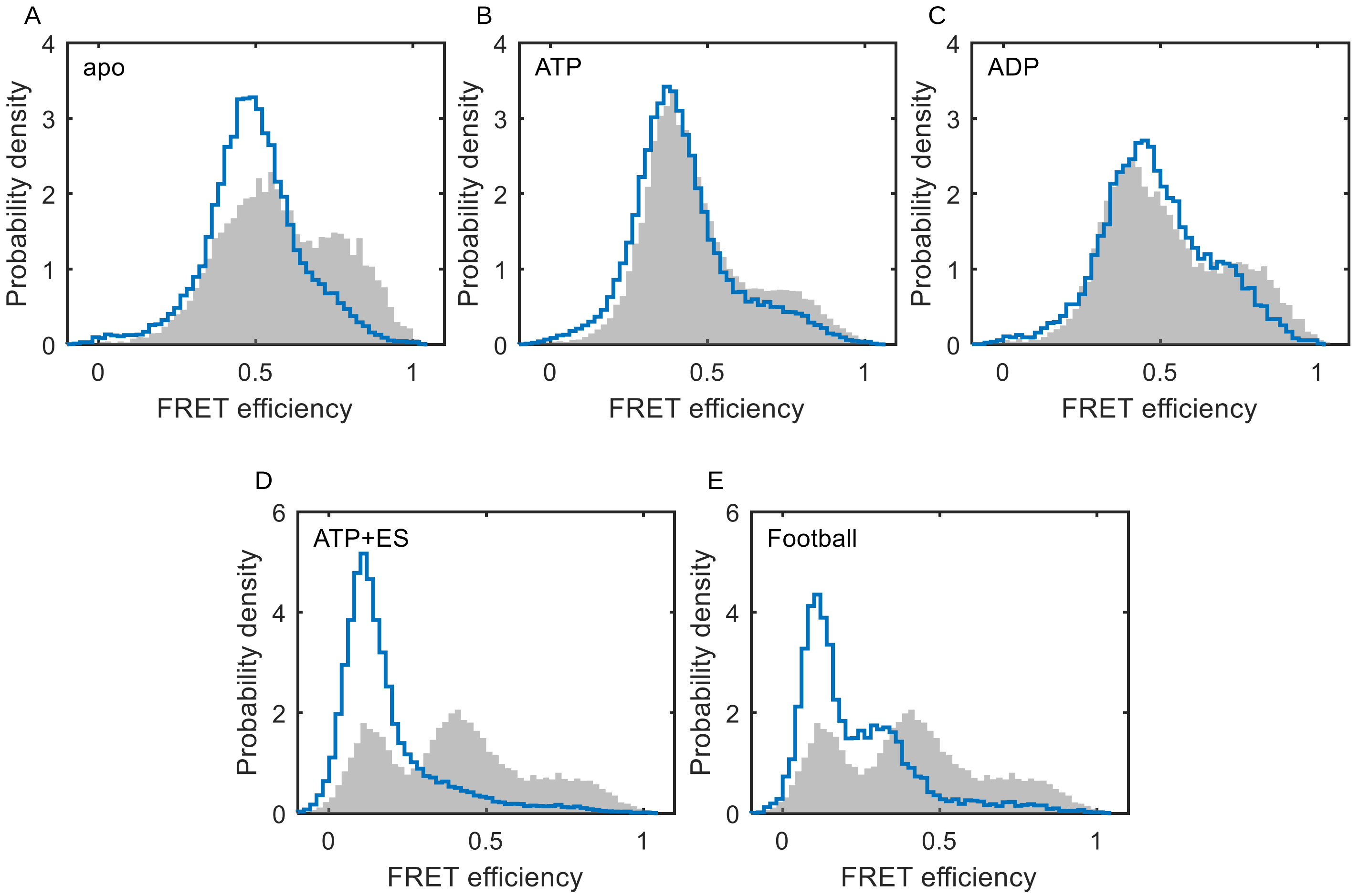


**Figure S5**: Comparison between corrected FRET efficiency histograms of GroEL 255C/428C double-ring (filled gray histograms) and SR1 (solid line histograms) constructs under different conditions: **A**) apo, **B**) 1 mM ATP, **C**) 1 mM ADP, and **D**) 1 mM ATP + ES. The FRET efficiency histograms are corrected for background, leaking, and direct acceptor excitation. All nucleotides and GroES (oligomer 100-500 nM) are in excess compared to GroEL (non-labeled + labeled oligomers 0.4-5 nM). The similar FRET efficiency manifolds suggest that the SR1 subunit adopts similar conformations as the double-ring subunit. Variations in the apo, ATP, and ADP conditions may originate because, during the smFRET diffusion experiment of the double-ring variant, the observed labeled subunit can be in the *cis* or the *trans* ring, giving rise to two indistinguishable types of conformational distributions in one experiment. This effect is best observed in the ATP-ES histogram (**D**), where FRET efficiency distributions of both low (ES-bound) and higher (nucleotide-bound) appear simultaneously. Conversely, the SR1 variant, having only one ring, adopts mainly a low FRET efficiency GroES-bound state. **E**) Comparison between FRET efficiency histograms of the double-ring complex in 1mM ATP+ES (filled histogram) and the symmetric GroEL football complex (solid line histogram). The football complex has two GroES oligomers bound to both GroEL rings, leading to an increase of the low FRET efficiency population in comparison to the asymmetric bullet complex.

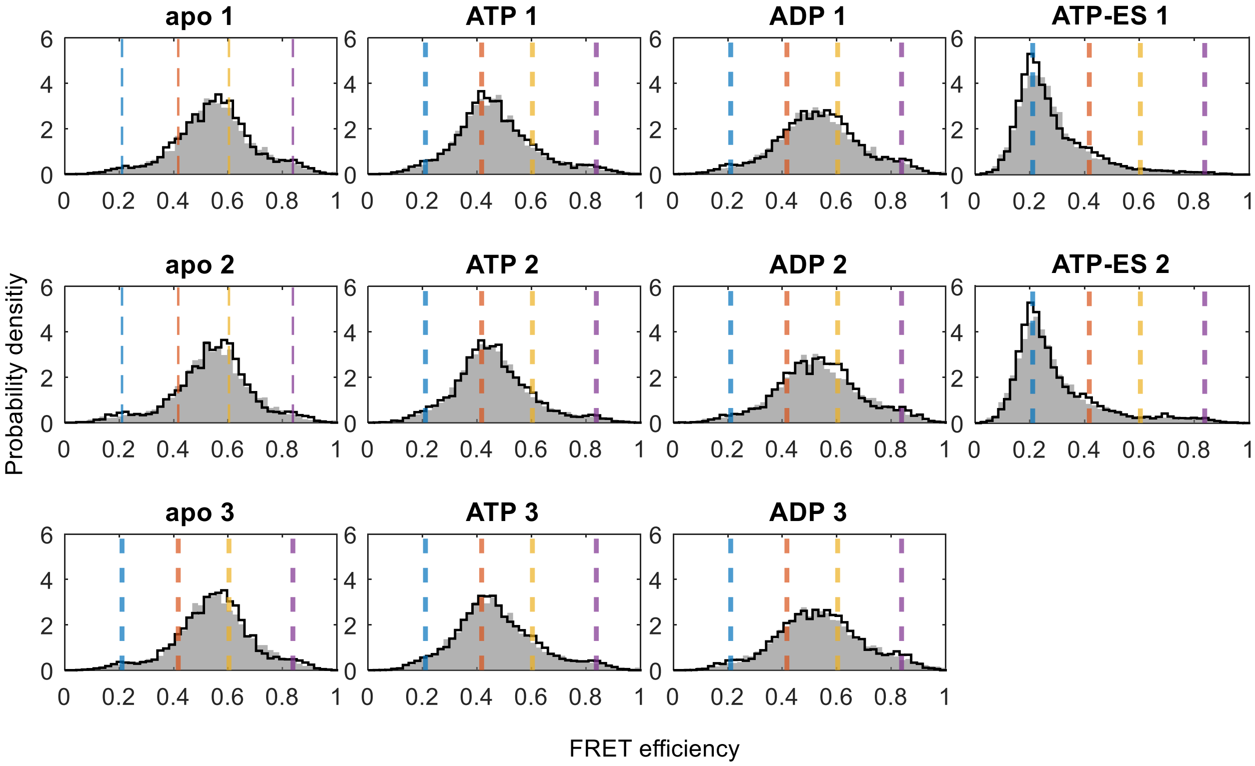


**Figure S6**: Raw FRET efficiency histograms of all measured datasets (gray) with recolored histogram (solid line) based on model parameters from the global $H^2MM$ analysis. The uncorrected FRET efficiency values of the four microstates are marked as color-coded dashed lines. The agreement between the raw and recolored FRET efficiency histograms demonstrates that the parameters found by the HMM global analysis successfully describe the experimental smFRET data.

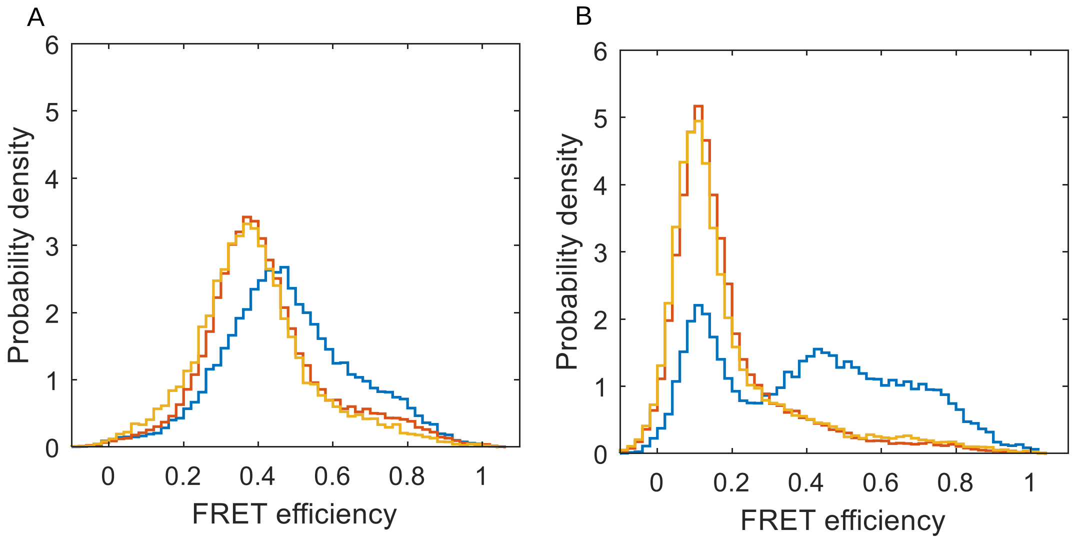


**Figure S7**: SR1 in the presence of the ATP analog ADP-BeFx. **A**) Corrected FRET efficiency histograms of SR1 in the presence of 1 mM ADP (blue), 1 mM ATP (red), and 1 mM ADP-BeFx (yellow). The similar FRET efficiency distributions in the presence of ATP and ADP-BeFx indicant that the γ-phosphate of ATP is essential for stabilizing the ATP-bound conformational state and that ADP hydrolysis is not the cause of the broad distribution. **B**) Corrected FRET efficiency histograms of SR1 in the presence of 1 mM ADP + GroES (blue), 1 mM ATP + GroES (red), and 1 mM ADP-BeFx + GroES (yellow). The GroES heptamer concertation is 150-300 nM. ADP allows a significant population of the low FRET efficiency conformational state, probably due to a lower affinity to GroES in the presence of this nucleotide. The similar FRET efficiency distributions in the presence of ATP and ADP-BeFx, compared to the ADP histogram, indicant that the γ-phosphate of ATP is essential for stabilizing the low FRET efficiency population of the GroES-bound conformational state.

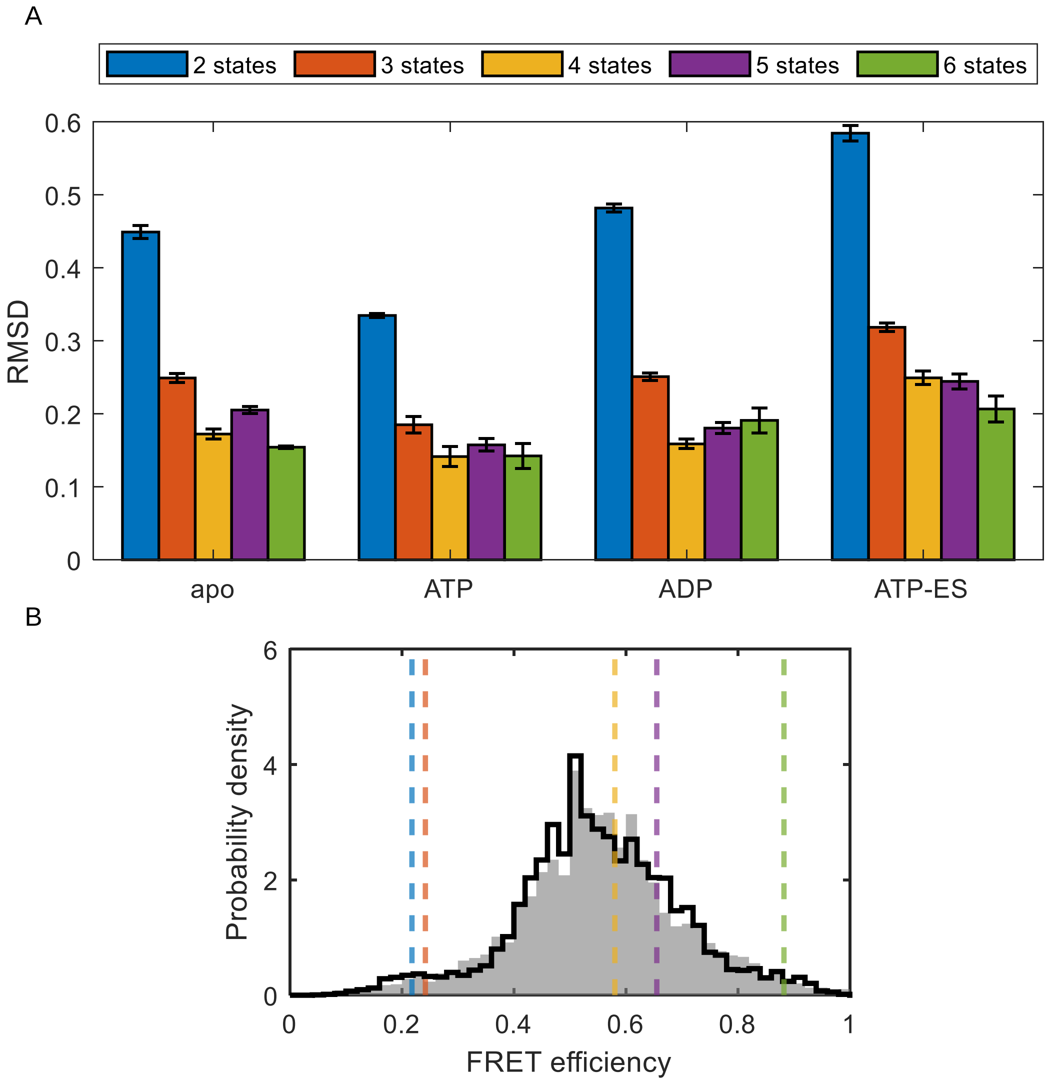


**Figure S8**: Test for model selection. **A)** Mean RMSD scores between experimental histograms and "recolored" FRET efficiency histograms. Shown are mean values and standard errors, based on three recoloring realizations. The lower the score, the better the match between the data and the $H^2MM$ model. As the number of states increases, so does the score. However, $H^2MM$ models with more than 4 states result in states with similar FRET efficiency values, implying redundancy. **B)** This panel demonstrates redundancy obtained in analysis 5 states. Notice that the two states with the lowest FRET efficiency values are essentially degenerate. The experimental and "recolored" histograms are in gray area and solid black line, respectively. The dashed lines show the positions of states obtained from the analysis.

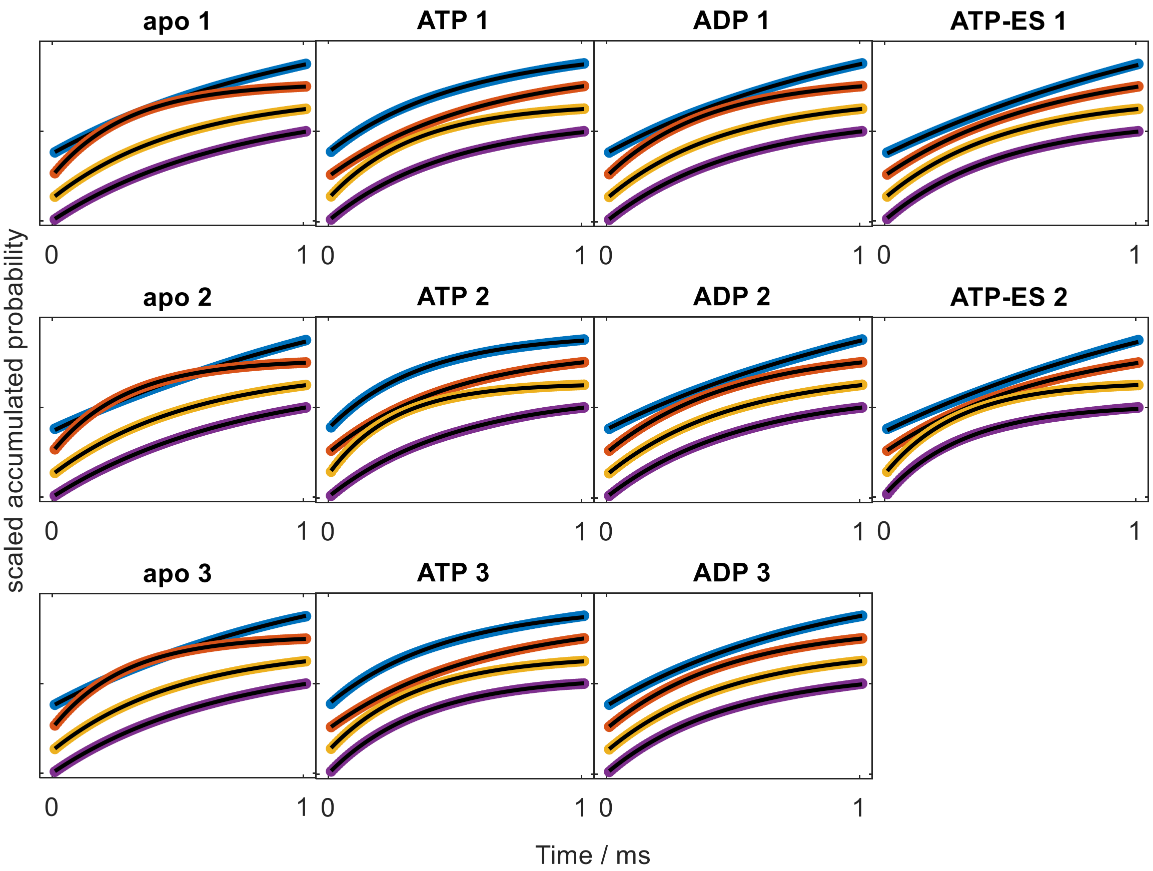


**Figure S9**: Likelihood-weighted dwell time analysis on datasets repeats. Dwell time distributions were generated for each of the datasets using the model parameters obtained in the $H^2MM$ analysis. Shown are the scaled accumulated dwell time distribution of each of the four states (same color code as elsewhere), each fitted to a mono-exponential function (solid black line). The plots are offset from each other for ease of observation. The resulting mean dwell time values from the fits, shown in **Table S2**, match the values from the $H^2MM$ parameters.

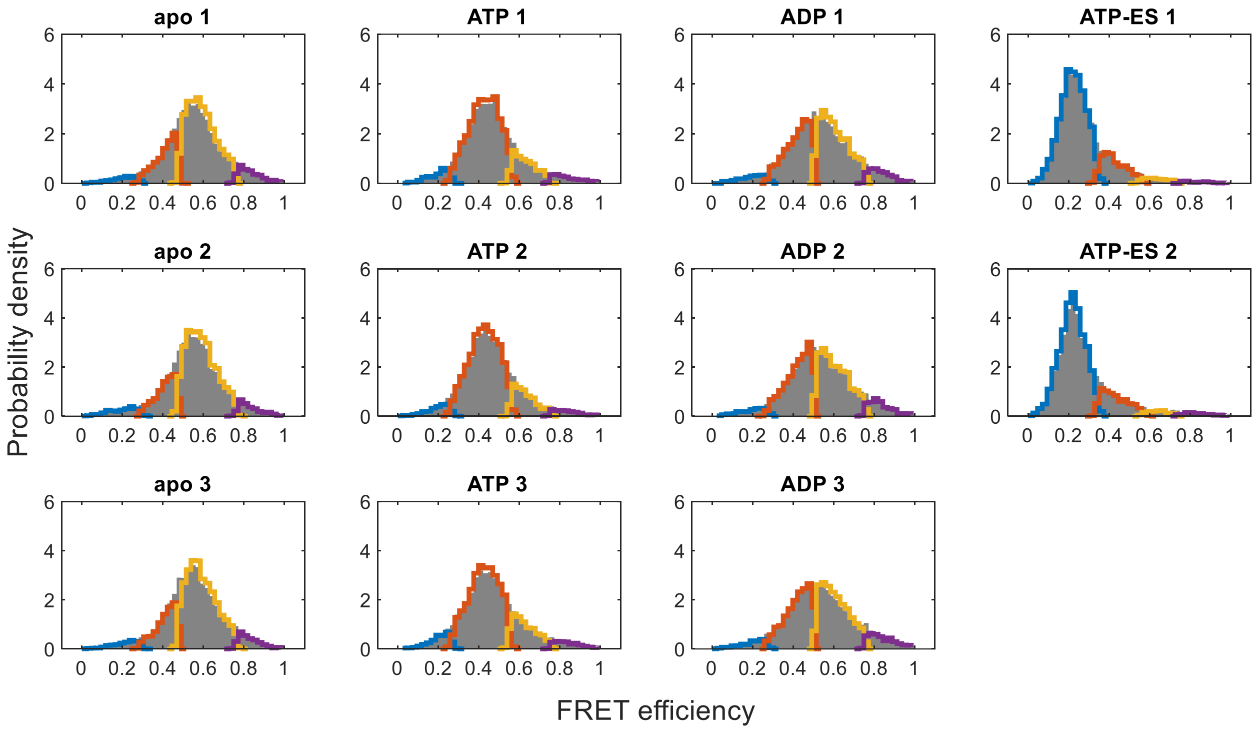


**Figure S10:** Raw FRET efficiency histograms of all GroEL datasets (gray) with individual microstate histograms based on Viterbi assignment, specified with the same color code used elsewhere in this work. The shown histograms were generated from photon trajectories that have no state-to-state transitions. The separation between the FRET efficiency distributions of the four microstates validates the ability of $H^2MM$ to correctly identify the microstates in the dataset.

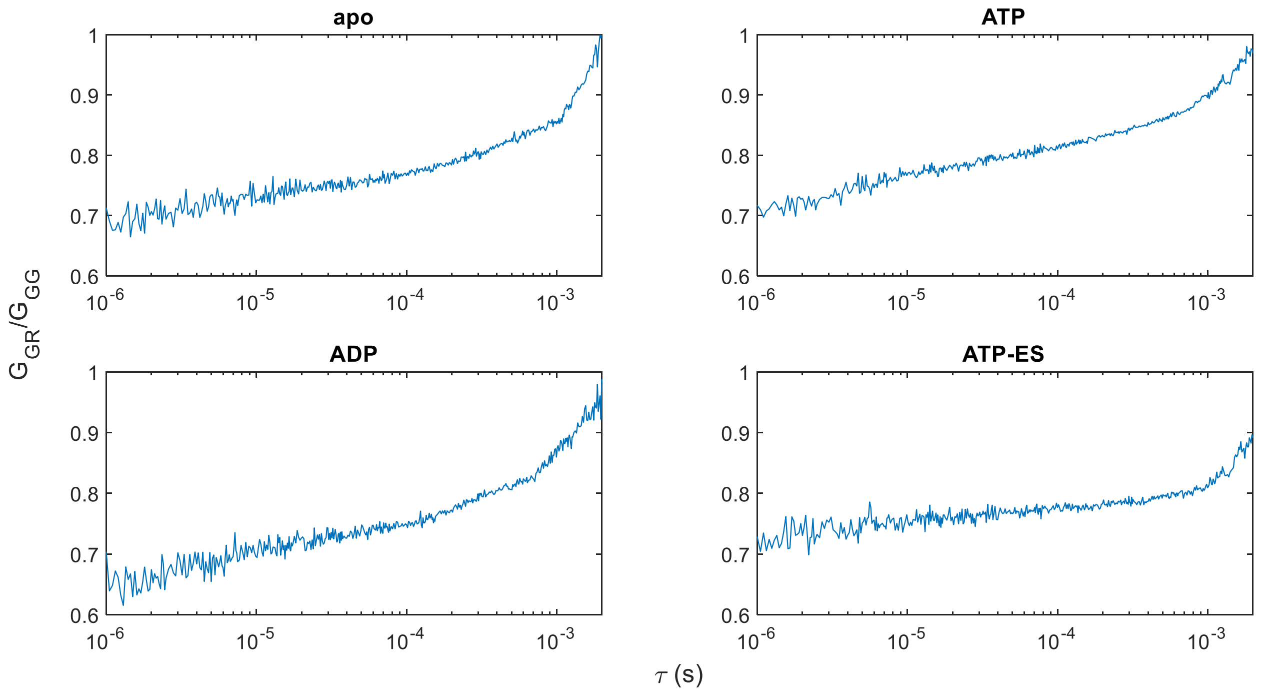


**Figure S11:** Burst-wise fluorescence correlation analysis demonstrates the existence of submillisecond conformational dynamics of the GroEL subunit under all measured conditions. Shown are the correlation ratios between the donor-acceptor cross-correlation function and the donor-donor autocorrelation function versus time shift (see Burst-Wise Correlation Analysis in Methods). This representation removes contributions due to diffusion. The correlation ratio plots show a bi-phasic increase: in the range of ~1-10 μs, presumably due to triplet-state dynamics and in the range of ~500-1000 μs due to conformational dynamics of the GroEL subunit. Notice the late increase in the case of ATP-ES, where the subunit is expected to show slower conformational dynamics.

### Supplementary tables

**Table S1**: Steady-state fluorescence anisotropy of SR1 labeled with a single dye.

| | apo | ATP | ATP ES |
|---|---|---|---|
| 255C Alexa488 | 0.20 ± 0.02 | 0.20 ± 0.01 | 0.18 ± 0.01 |
| 255C Alexa594 | 0.27 ± 0.01 | 0.25 ± 0.01 | 0.25 ± 0.01 |
| 428C Alexa488 | 0.192 ± 0.002 | 0.173 ± 0.004 | 0.170 ± 0.001 |
| 428C Alexa594 | 0.21 ± 0.01 | 0.216 ± 0.004 | 0.204 ± 0.002 |

**Table S2:** Dwell time values for each microstate under different conditions obtained from $H^2MM$ parameters, and weighted dwell-time analysis (DTA) fits (see **Figure S9**). The values are in milliseconds and presented as mean with standard error from 2-3 repeats.

| Condition | Microstate 1 | | Microstate 2 | | Microstate 3 | | Microstate 4 | |
|---|---|---|---|---|---|---|---|---|
| | $H^2MM$ | DTA | $H^2MM$ | DTA | $H^2MM$ | DTA | $H^2MM$ | DTA |
| Apo | 0.65 ± 0.03 | 0.72 ± 0.02 | 0.49 ± 0.01 | 0.53 ± 0.01 | 0.284 ± 0.003 | 0.296 ± 0.003 | 1.3 ± 0.2 | 1.2 ± 0.1 |
| ATP | 0.39 ± 0.02 | 0.46 ± 0.03 | 0.29 ± 0.02 | 0.32 ± 0.02 | 0.55 ± 0.03 | 0.56 ± 0.03 | 0.37 ± 0.03 | 0.46 ± 0.04 |
| ADP | 0.45 ± 0.01 | 0.51 ± 0.01 | 0.451 ± 0.007 | 0.481 ± 0.007 | 0.45 ± 0.03 | 0.46 ± 0.03 | 0.91 ± 0.08 | 1.1 ± 0.1 |
| ATP+ES | 0.33 ± 0.05 | 0.40 ± 0.05 | 0.35 ± 0.06 | 0.39 ± 0.06 | 0.616 ± 0.009 | 0.665 ± 0.002 | 2.0 ± 0.1 | 1.459 ± 0.001 |

**Table S3:** Populations of microstate presented in %.

| Macrostate | Microstate 1 | Microstate 2 | Microstate 3 | Microstate 4 |
|---|---|---|---|---|
| apo | 8.4 ± 0.5 | 55.1 ± 0.1 | 29.6 ± 0.6 | 7.0 ± 1.0 |
| ATP | 4.90 ± 0.05 | 25.3 ± 0.7 | 58.4 ± 0.7 | 11.5 ± 0.7 |
| ADP | 9.4 ± 0.2 | 44.6 ± 0.7 | 38.4 ± 0.9 | 7.6 ± 0.6 |
| ATP-ES | 3.2 ± 0.5 | 6.23 ± 0.06 | 25.4 ± 1.0 | 65.2 ± 0.4 |
